# Genome size evolution and phenotypic correlates in the poison frog family Dendrobatidae

**DOI:** 10.1101/2023.06.30.547273

**Authors:** Tyler E Douglas, Roberto Marquez, V. Renee D Holmes, J. Spencer Johnston, Rebecca D Tarvin

## Abstract

Adaptive and neutral processes have produced a spectrum of genome sizes across organisms. Amphibians in particular possess a wide range in C-values, from <1 to over 125 pg. However, the genome size of most amphibians is unknown, and no single family has been comprehensively assessed. We provide new estimates for 32 poison frog species representing the major lineages within Dendrobatidae using Feulgen staining of museum specimens and flow cytometry of fresh tissue. We show that genome size in Dendrobatidae has likely evolved gradually, with potential evolutionary rate shifts in the genera *Phyllobates* and *Hyloxalus*, which respectively possess species with the largest (13.0 pg) and second smallest (2.6 pg) genomes in the family. Phylogenetically controlled regression analyses indicate that genome size is positively correlated with snout-vent-length, oocyte number, and clutch size, but negatively correlated with active metabolic rate and metabolic scope. While body size and metabolic rate are also correlates of toxicity, we found no relationship between genome size and evolution of chemical defense within Dendrobatidae. Genome size evolution in Dendrobatidae provides insight into the processes shaping genome size evolution over short timescales and establishes a novel system in which to study the mechanistic links between genome size and organismal physiology.

## Resumen

Los procesos adaptativos y neutrales han producido un espectro de tamaños de genoma a través de la diversidad biológica. Los anfibios en particular poseen una amplia gama de valores C, desde <1 pg hasta más de 125 pg. Sin embargo, se desconoce el tamaño del genoma de la mayoría de especies de anfibio, y no se ha evaluado exhaustivamente su evolución en detalle dentro de ninguna familia. Proporcionamos estimados del tamaño del genoma para 32 especies de ranas venenosas que representan los principales linajes de la familia Dendrobatidae utilizando la tinción de Feulgen con especímenes de museo y citometría de flujo con tejido fresco. Mostramos que el tamaño del genoma en Dendrobatidae probablemente evolucione bajo el fecto de selección purificadora, potencialmente con regímenes de selección distintos al ancestral géneros *Phyllobates* e *Hyloxalus*, que respectivamente poseen especies con el genoma más grande (13,0 pg) y el segundo más pequeño (2,6 pg) de la familia. Análisis de regresión con control filogenético indican que el tamaño del genoma se correlaciona positivamente con la longitud hocico-cloaca, el número de oocitos y el tamaño de la nidada, y negativamente con la tasa metabólica activa y el alcance metabólico. Aunque el tamaño corporal y la tasa metabólica se correlacionan con la toxicidad, no encontramos relación entre el tamaño del genoma y la evolución de la defensa química dentro de Dendrobatidae. Nuestra evaluación del tamaño del genoma en Dendrobatidae proporciona información sobre los procesos que dirigen la evolución del tamaño del genoma en escalas de tiempo cortas y un nuevo sistema que facilitará las investigaciones de los vínculos mecánicos entre el tamaño del genoma y la fisiología del organismo.

## Introduction

Genomes can expand or contract through nonadaptive and adaptive mechanisms. In eukaryotes, genomes expand primarily through the self-replication of transposable elements (TEs), which generates long stretches of repetitive DNA (Kidwell 2002; Sun et al. 2012). Genomic TE activity largely explains why genome size does not scale with gene number in eukaryotes (the C-value paradox; Cavalier-Smith 1978). Genome expansion may be deleterious because accumulating introns and repetitive elements could increase mutational load and mutational hazard (Lynch et al. 2006), and set lower limits to traits like cell size and the rate of cell replication (Gregory 2002). Under this view, genome expansion is a function of genetic drift, and genome expansion is likely in populations where purifying selection is weak and genetic drift is strong (i.e., small effective population size Ne; Lynch and Conery 2003). Notably, however, some classical studies of neutral genome size evolution included data spanning prokaryotes and eukaryotes without phylogenetic correction and may have overestimated the role of neutral evolution in generating variation in genome size (Whitney and Garland 2010; Smith 2016).

Several selective hypotheses for the evolution of genome size exist based on observed correlations between genome size and fitness-related traits. For example, in some species genome size correlates positively with cell size and body size and negatively with developmental rate, developmental complexity (extent of tissue specialization), and metabolic rate (Cavalier-Smith 1982; Jockusch 1997; Kozłowski et al. 2003; Womack et al. 2019). Ecological or physiological constraints on any of these traits could indirectly impose selection on genome size. For instance, miniaturization is associated with genome size reduction, which results in smaller cells and makes achieving the minimum number of cells required to build functional organs possible (Levy and Heald 2015; Decena-Segarra et al. 2020). On the other extreme, large genomes may require physiological accommodations, such as enucleation of red blood cells in plethodontid salamanders with especially large genomes (Mueller et al. 2008). However, the direction and magnitude of physiological correlates with genome size vary between taxonomic groups. For instance, metabolic rate correlates negatively with genome size in birds, but this correlation is only weakly supported in mammals and amphibians (Licht and Lowcock 1991; Vinogradov 1995; Gregory 2002). Thus, the adaptive dynamics underlying genome size evolution may be lineage-specific and/or influenced by multiple independent traits, and therefore difficult to discern without denser taxon sampling at lower taxonomic levels.

Amphibians in particular are an ideal group for testing evolutionary hypotheses of genome size evolution, as they vary dramatically in genome size (Liedtke et al. 2018) and several of its potential life history correlates, including body size, cell size, ontogeny, reproductive ecology, and metabolic rates (Gomes et al. 2004; Elinson and del Pino 2012; Levy and Heald 2015; Liedtke et al. 2022; Johnson et al. 2023). Within amphibians, three clades of the family Dendrobatidae (*sensu* Santos et al. 2009) have independently evolved bright coloration and the ability to sequester dietary toxins, along with other correlated changes, together known as the aposematic syndrome (Santos et al. 2003; Vences et al. 2003; Santos and Cannatella 2011). Chemical defense evolution has been associated with higher aerobic capacity (Santos and Cannatella 2011) and substantial variation in body size (Silverstone 1975, 1976; Santos and Cannatella 2011), which, in some chemically defended species, positively correlates with defensive alkaloid quantity (Saporito et al. 2010). Furthermore, *HSP90*, a heat-shock protein that could be involved in suppressing transposable elements in multiple animal lineages, such as *Drosophila* (Specchia et al. 2010; Gangaraju et al. 2011), rodents (Olivieri et al. 2012; Xiol et al. 2012), and nematodes (Ryan et al. 2016), was shown to bind to alkaloids (Caty et al. 2019). Thus, natural selection for chemical defenses could indirectly impact genome size by acting on body size, metabolic rate, or *HSP90* function. Because the effects of body size and metabolic rate on genome size are thought to operate in competing directions, the specific effects that chemical defense would have on genome size are unclear. Here, we leverage the variation of a suite of traits in Dendrobatidae (Santos et al. 2003) to investigate the patterns and possible drivers of genome size evolution. Taking advantage of the diversity of preserved specimens available in natural history collections, we generated genome size estimates for 32 species, and used phylogenetic comparative methods and publicly available phenotypic data to inquire into the drivers of genome size evolution in dendrobatids.

## Methods

### Specimen identification

Species identities of museum specimens and data in the Animal Genome Size Database (AGSD) were checked using locality information and photographs or specimens when available (Table S1; see supplementary materials for full details).

### Cell extraction, staining, and imaging

We extracted red blood cells from formalin-fixed specimens stored in ethanol in the Museum of Vertebrate Zoology (MVZ). Our general approach follows Hardie et al. (2002), with modifications by Womack et al. (2019), who specifically analyzed formalin-fixed amphibian specimens. Cells were stained and imaged as detailed below in two separate batches. Batch one included 22 dendrobatid species and 6 non-dendrobatid species with published genome sizes; batch two included 10 dendrobatid species and 12 non-dendrobatid species (Table S2). We drew blood cells 4 times from 1-2 individuals per species, deposited cells arbitrarily across a set of microscope slides to prevent batch bias and allowed the cell extracts to air dry for 30 minutes. We then used Feulgen staining as detailed in Hardie et al. (2002). In brief, we fixed cells in 85 methanol:10 formalin:5 acetic acid for 24 hours, rinsed cells under running water for 10-minutes, hydrolyzed cells in 5.0 N HCl for 2 hours, briefly submerged cells in 0.1 N HCl, then stained cells for 2 hours with Schiff’s reagent (Fisher Scientific 632-99-5). Next, we rinsed cells with bisulfite solution 3 times for 5 minutes, followed by a 10-minute wash under running water and 3 2-minute rinses in distilled water. We air-dried slides and mounted cover glass on slides using Permount (Fisher SP15500).

We imaged cells using a SeBa camera (SEBACAM5C) mounted to an Laxco LMC-2000 microscope with a 100x immersion oil lens. We white-balanced each slide individually using SeBa imaging software SebaView v3.7 prior to imaging.

### Genome and cell size estimation

Using ImageJ v1.53c we calculated the integrated optical density (IOD) from cell images, which measures the amount of stained material (i.e., DNA), in 6-66 nuclei per species (average = 19.93 nuclei, SD = 12.18). To convert IOD values to genome size estimates, we generated IOD measurements for 15 species with published genome sizes and 2 species for which we estimated genome sizes using flow cytometry (Tables S1 and S2; see below for methods), and used them to generate a standard curve. When possible we selected species with multiple published genome size measurements for use in the standard curve. We obtained the median IOD across nuclei for each specimen, as IOD scores exhibited left or right-skewed distributions in some samples. For species with more than one sampled specimen we obtained the median of specimen medians.We built our standard curve as a linear mixed-effects model with genome size as the dependent variable, IOD as a fixed-effect predictor, and sampling batch as a random-effect predictor, using the lmer() function in the *lme4* R package (Bates et al. 2015).

The use of cells from formalin-fixed specimens for genome size estimation based on Feulgen staining can result in artifacts because Schiff’s reagent binds aldehydes (including formaldehyde), and preservation and long-term storage can result in cell damage (Greilhuber and Temsch 2001; Hardie et al. 2002). This being said, multiple studies have quantified genome size and other morphonuclear parameters using Feulgen-stained nuclei obtained from formalin-fixed specimens, both from natural history (e.g., Itgen et al. 2019, 2022; Womack et al. 2019; Wang et al. 2021), and hospital archival collections (e.g., Dorman et al. 1990; Salmon et al. 1992; Fang et al. 2004), with accuracy often similar to other methods including flow cytometry. At least in some cases, such measurements have not been found to be particularly susceptible to variation in the formalin fixation process and posterior storage conditions (Schimmelpenning et al. 1990), although more general tests of these remain scarce. With this in mind, Feulgen-based genome size estimates from formalin-fixed specimens can be considered reliable, as long as the quality of such measurements is first evaluated, for instance by the inclusion of a set of samples whose parameters of interest are known (Greilhuber and Temsch 2001; Greilhuber 2008). In the case of amphibians, studies using Feulgen staining on formalin-fixed cells have obtained tight-fitting correlations between IOD and genome size determined using flow cytometry (e.g., Itgen et al. 2019; Womack et al. 2019), R^2^ = 0.91-0.93), close to those from studies using alcohol-preserved (e.g. Liedtke et al.2018, R^2^ = 0.99) or fresh (e.g., Decena-Segarra et al. 2020)^1^) tissues.

We ensured the reliability of our estimates in three complementary ways: First, we removed specimens of *Pseudacris nigrita*, *Oophaga pumilio*, and *Phyllobates terribilis* from the calibration curve because cells from these specimens showed clear morphological anomalies. Second, we evaluated the goodness of fit of our calibration curve using Nakagawa and Schielzeth’s (2013) marginal R^2^, which quantifies the variance explained by the fixed effects of a mixed-effects model, using the r.squaredGLMM()function of the *MuMIn* R package (Bartoń 2024), and generated confidence intervals around our calibration curve parameters via bootstrapping to evaluate its precision and susceptibility to outlier specimens. Finally, we included seven species in both staining batches to assess repeatability in genome size estimates between batches.

To estimate the cell size of each species we manually traced the cell membrane of 2-22 cells per species (average = 12.30 cells, SD = 5.07) in ImageJ and calculated the area within each cell. Cell sizes were averaged by species for subsequent analysis.

### Flow cytometry

We used flow cytometry to generate genome size estimates for four species of the genus *Phyllobates*: *P. terribilis, P. bicolor, P, aurotaenia*, and *P. vittatus*. All frog-rearing protocols were approved by the University of Michigan’s Institutional Animal Care and Use Committee (protocol #PRO00010325). Captive-bred specimens were euthanized with an overdose of topical benzocaine, and muscle samples were immediately dissected and flash-frozen on dry ice, and kept frozen until further use. See above for animal care and use protocol details. Genome size was estimated as described by Johnston et al. (2019) and details of flow cytometry methods are located in supplementary methods.

### Phylogenetic inference

To investigate the evolutionary patterns and correlates of genome size from a phylogenetic perspective, we generated a time-calibrated phylogeny of our study species based on the relationships inferred by Grant et al. (2017) and Vacher et al. (2017), using publicly available DNA sequences from six loci (one mitochondrial, five nuclear; Table S3). We pruned the Grant et al. (2017) tree to include only our target species using *ape* (in R v5.3; Paradis and Schliep 2019)*. Anomaloglossus surinamensis* was not included in the Grant et al. (2017) phylogeny, so we placed it as the sister species to *Anomaloglossus stepheni* following Vacher et al. (2017). Although *Aromobates tokuko* was recently shown to have a remarkably small genome (0.8 pg; Liedtke et al. 2018), there are no DNA sequences available for this species, so we did not include it in our analyses. Because dendrobatid systematics remain understudied, and there is evidence for both cryptic species and species complexes (e.g., Tarvin et al. 2017; Guillory et al. 2019; Márquez et al. 2020), we used tree tips and sequences from the closest available locality to the collection site of the specimens used to estimate genome size (details in Table S3).

We aligned sequences for each locus (Table S3) using MUSCLE v3.8.1551 (Edgar 2004), and chose a partitioning strategy and models of sequence evolution using PartitionFinder2 (Lanfear et al. 2017). The final (concatenated) matrix contained 5,493 bp (2,513 bp from mtDNA; 2,980 bp across 5 nuclear loci ranging between 316 bp to1,200 bp), with 35% gaps and 1204 parsimony-informative sites. We then ran a concatenated analysis on BEAST 2.6.5 (Bouckaert et al. 2019), setting the Grant et al. (2017) topology as a hard constraint, and using three normally-distributed calibration priors based on previous studies: The root of Dendrobatidae (i.e. the root of the tree) was set to N(μ = 38.534, σ = 5.151) following Santos et al. (2009), and the nodes corresponding to Dendrobatinae and Colostethinae (*sensu* Grant et al. 2006) were set to N(μ=26.655, σ=4.791) and N(μ=19.392, σ=3.793), respectively, based on Guillory et al. (2019).

We used empirical base frequencies and a calibrated Yule tree prior (Heled and Drummond 2012), and set the Yule birth rate and clock rate priors to gamma(0.01, 50), and the GTR relative rate priors to *gamma*(2, 0.5) for transitions and *gamma*(2, 0.25) for transversions. All other priors were left at the default. Independent uncorrelated log-normal clocks were set for all partitions, and a preliminary MCMC was run for 5 million steps, sampling every 5,000. We recovered posterior distributions of the parameters describing clock rate variation among branches (*ucldStdev*) that leaned heavily towards zero in four of the five nuclear loci, suggesting very little rate variation at these genes, so they were modeled under a strict clock in further runs. To obtain a final tree, we ran two independent MCMC samplers for 20 million steps, sampling every 5,000, and merged their results after discarding the initial 10% of each one as burnin. Convergence and mixing within and between runs were assessed based on effective sample sizes (ESS) estimates and trace plots generated in Tracer v1.7.1 (Rambaut et al. 2018). A maximum clade credibility (MCC) tree with common ancestor node heights was produced in TreeAnnotator (distributed with BEAST) and used for all downstream analyses. The xml file used as input for BEAST2, which includes alignments and software parameters, is available on the Dryad repository for this paper.

### Patterns of genome size evolution

We explored the patterns of genome size evolution along the dendrobatid phylogeny in several complementary ways. First, we used BAMM (Rabosky 2014) to estimate its rate of evolution across the tree, and to assess the degree to which discrete rate shifts at specific branches can explain the extant variation in dendrobatid genome size. We used the R package BAMMtools (Rabosky 2014) to generate priors appropriate for the temporal and taxonomic scale of our tree and phenotype, and ran BAMM for 10,000,000 steps, sampling every 5,000, for a total of 20,000 samples from the posterior, out of which we discarded 10% as burnin. From the post-burnin BAMM samples, we estimated the posterior distribution for the number of rate shifts along the tree, as well as the frequency of different rate shift configurations and their associated rates of phenotypic evolution using BAMMtools.

Next, we examined the evidence for discrete (i.e., saltational) evolutionary regime shifts in the evolution of genome size using the *l1ou* R package (Khabbazian et al. 2016) to fit and rank all possible single and multi-optimum Ornstein-Uhlenbeck (OU) models of phenotypic evolution (Hansen 1997) for our tree. Briefly, these models allow different subtrees to have distinct evolutionary “optima,” which we can use to gauge whether specific branches of the tree have experienced rapid jumps in genome size. We used the default lasso approach implemented in *l1ou*, and model fit was assessed using the phylogenetically corrected Bayesian information criterion (pBIC; Khabbazian et al. 2016). To quantify the strength of support for each model, we calculated the pBIC weight (*w_i_* for model *i*) as

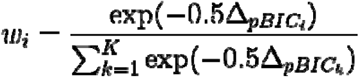

Where Δ*_pBICi_* is the difference between a given model (model *i*) and the best model (i.e., the one with the smallest pBIC), and *K* is the total number of models.

Finally, we evaluated the fit of different continuous trait evolution models to our data, in order to gain further insight into the processes driving the evolution of genome size in dendrobatids. The analyses above suggested there may have been saltational events in the evolution of genome size (see Results section), so we first selected a set of four single and multi-regime model configurations, and fit Brownian Motion (Felsenstein 1973), and OU models based on these regimes. In addition, we fit Early Burst (Harmon et al. 2010) and White Noise models across the whole tree. BM models were fit using the brownie.lite() function in *phytools* (Revell 2024), OU models were fit using the fit_OU() function in *l1ou*, and the EB and White Noise models were fit using the fitContinuous() function of the *geiger* package (Pennell et al. 2014). Model fit was evaluated using the corrected Akaike Information Criterion (AICc), since pBIC has not been implemented beyond OU models. To ensure compatibility between model-fitting algorithms, we also fit the single-regime BM and OU models with fitContinuous(), which obtained identical likelihoods and parameter estimates to the functions mentioned above.

### Correlates of genome size

To assess phenotypic correlates of genome size, we obtained data for several physiological, ecological, and life history traits from compilations available in the literature (Santos and Cannatella 2011; Santos 2012; Carvajal-Castro et al. 2021; full list of traits available in Table 1). In addition, we obtained elevation from the collection localities of the specific museum specimens used for genome size estimation when available, or the literature otherwise. We used snout-vent length (SVL) as a proxy for body size, which we measured directly on adult specimens or their photographs, except for the four *Phyllobates* species used in flow cytometry, for which SVL values were pulled from the literature (Márquez et al. 2020), as study specimens were juveniles. Finally, we scored each species as able or unable to sequester alkaloids based on data compiled by Santos, Tarvin, and O’Connell (2016). All trait data are available in Table S4; note that up to 47% of species are missing data for some traits, so the results of our correlations should be taken as preliminary evidence supporting hypotheses that require further validation.

To assess trait correlations, we fit univariate phylogenetic-generalized-least-squares (PGLS; Grafen 1989; Martins and Hansen 1997) linear regressions using the phylolm()function of the *phylolm* R package (Ho and Ané 2014). Covariance between traits due to phylogeny was modeled under the OU model with an estimated ancestral state (OUfixedRoot model). We added mass as a covariate in models involving metabolic rates (data were obtained from Santos 2012). To visualize relationships between traits as well as their phylogenetic distribution we generated phylomorphospace plots (Sidlauskas 2008) using *phytools*. Because some species of *Phyllobates* were outliers in terms of genome size and some of the tested predictors (see Results), and may therefore generate overestimates of phenotypic correlations, we ran our linear regressions both including and excluding this genus from the dataset.

## Results

### Genome size estimation

Our calibration curve displayed a very close fit between IOD and published genome size for all species in both staining batches (marginal R^2^ = 0.94), with narrow conference intervals around the slope and intercept (Fig. 1A, B), and highly consistent estimates between batches (Fig. 1C). In light of these results, our Feulgen-based estimates using formalin-fixed cells can be considered good proxies for genome size to evaluate the general patterns and correlates of this trait across the range of dendrobatid genome sizes. We note, however, that for more precise purposes, such as validating the completeness of *de novo* genome assemblies, or for intraspecific analyses, the Feulgen-based genome size estimates presented here (and in other similar studies) should be verified using additional methods, such as flow cytometry. Based on our IOD curve and flow cytometry workflow, we estimated the genome size of 32 species with previously unknown genome sizes (Table S5). Our estimated genome sizes (C-value) in Dendrobatidae ranged from 2.6 pg (*Hyloxalus infraguttatus*) to 13.0 pg (*Phyllobates bicolor*; Fig. 2).

**Figure 1.**
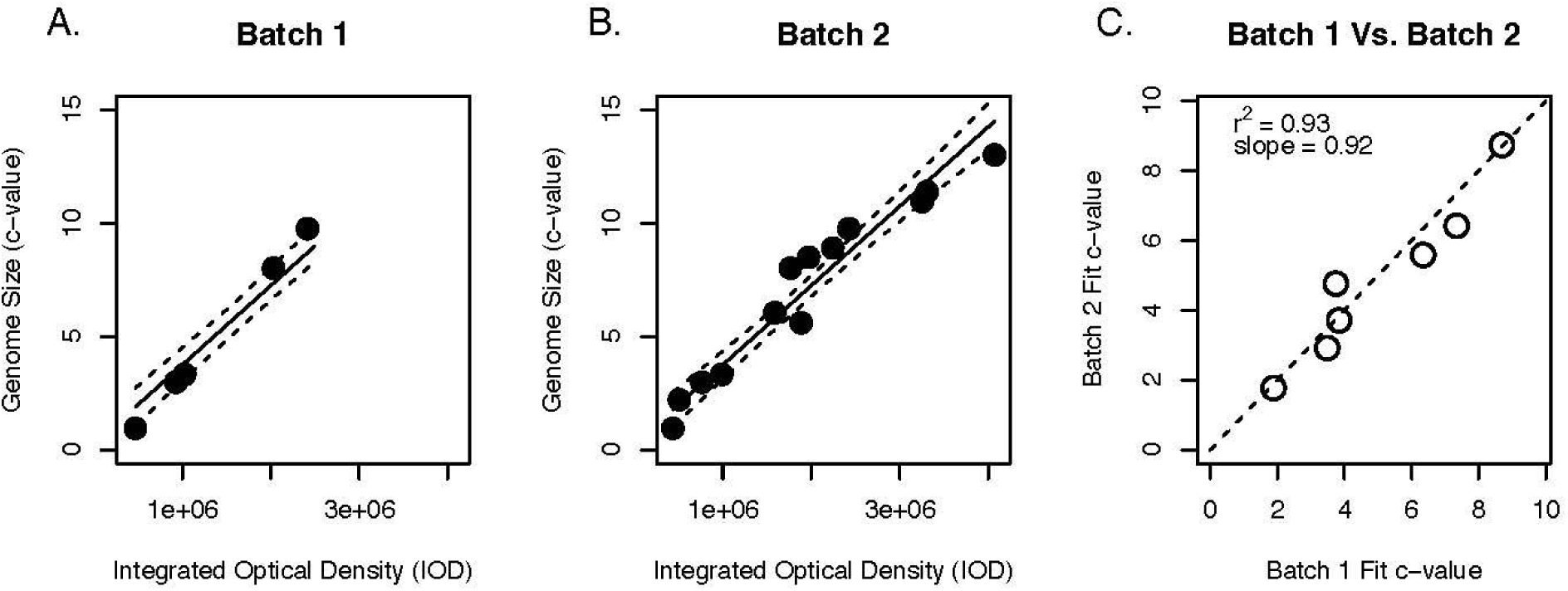
Standard curves used to convert IOD values to genome sizes. Panels A and B represent the curves inferred for each independent imaging batch. Solid lines represent the slope and intercept for each specific batch and broken lines 95% confidence intervals. Panel C shows the repeatability of our estimates by plotting genome size estimates for the seven species that were imaged in both batches. The diagonal represents the identity line.

**Figure 2.**
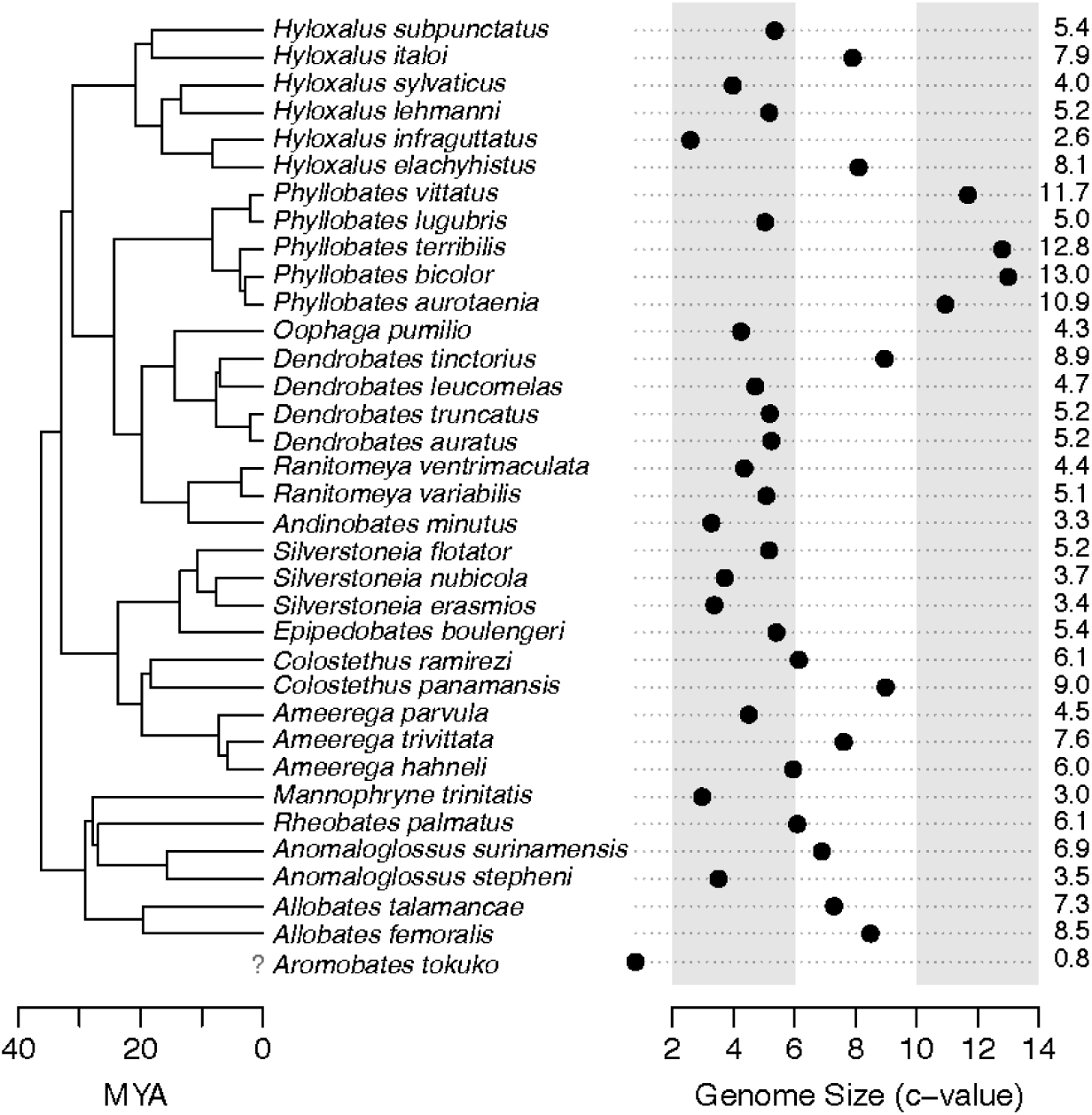
Genome size variation across the dendrobatid phylogeny. The position of *A. tokuko* is uncertain. Numbers to the right of the dot plot represent estimated or measured genome sizes for each species in pg. MYA: Million Years Ago.

### Patterns of genome size evolution

The rate of genome size evolution was relatively similar across the dendrobatid tree and stable through time, with faster rates concentrated in branches containing the five species of *Phyllobates* and *Hyloxalus elachyhistus* and *H. infraguttatus* (Fig. 3AB). Rate estimates from BAMM were generally concordant with those estimated under Brownian Motion (the BAMM phenotypic model assumes Brownian evolution; Rabosky et al. 2013), suggesting these estimates are accurate despite the number of species analyzed (Fig. S1). BAMM and *l1ou* both provided some evidence for discrete rate shifts. Specifically, the posterior distribution for the number of rate shifts recovered by BAMM peaked at 1-2 shifts (Fig. 3C), and the top models in all analyses contained scenarios with 2-3 evolutionary regimes that were better or comparable with single-regime models, although single-regime scenarios were also well supported (Fig. S2).

**Figure 3.**
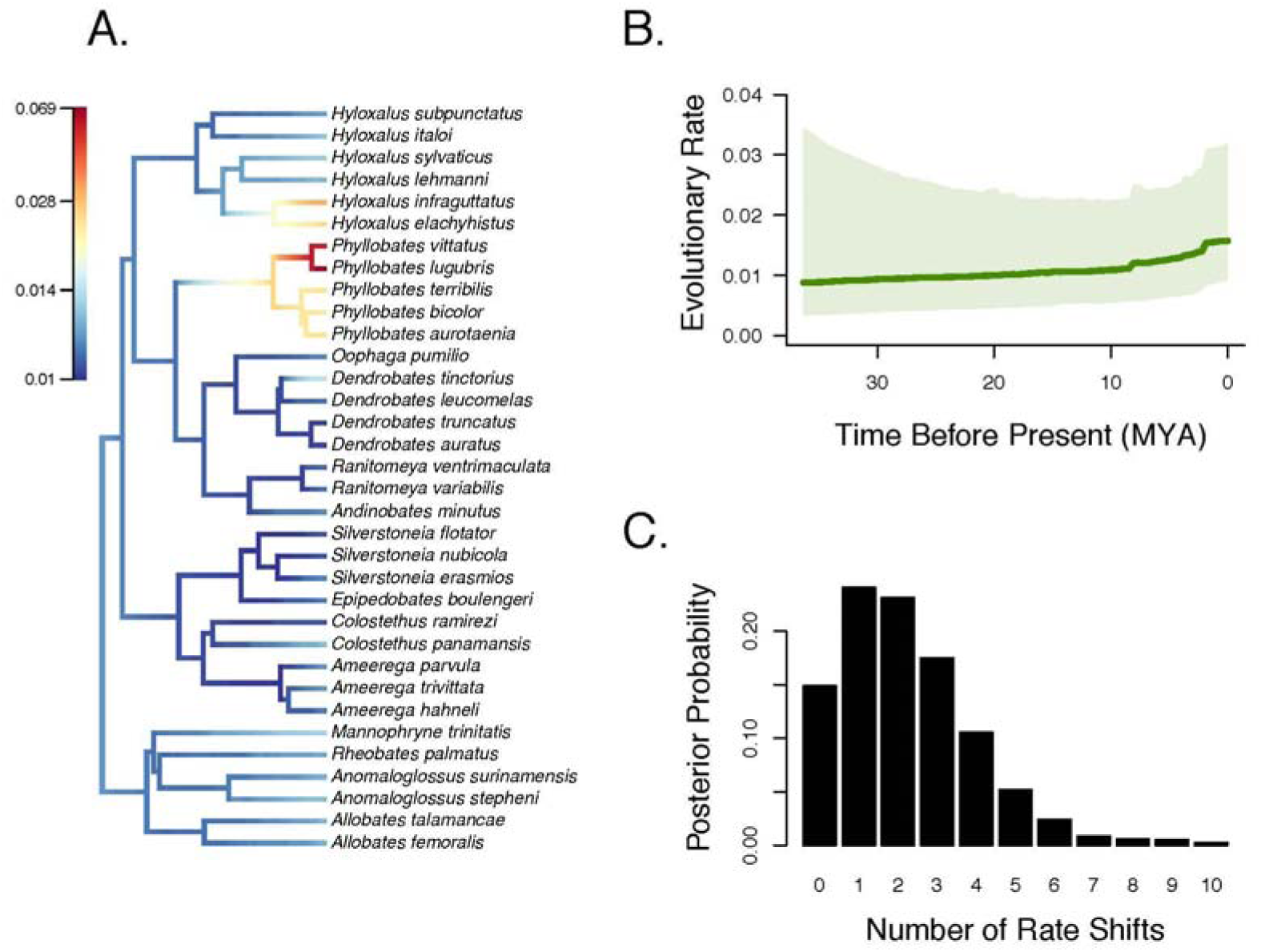
Rates of genome size evolution in dendrobatid frogs. A) Instantaneous rates of phenotypic evolution along branches of the dendrobatid phylogeny and B) across the phylogeny over time. C) Posterior distribution of the number of discrete rate shifts recovered by BAMM. The color ramp in panel A was generated using the Jenks method in the plot.bammdata() function of BAMMtools. Panel B is based on 100 time slices, the central line depicts the median posterior rate, and the surrounding polygon the 95% highest posterior density.

Based on the above evidence for multiple regimes of genome size evolution in different branches of the dendrobatid tree, we conducted a model-fitting exercise where we compared ten different single and multi-regime models of trait evolution, evolving under BM, OU, and EB dynamics (Fig. 3). The two best-fitting models, which together accounted for 74% of AICc weight were both multi-regime OU models with either *Phyllobates* departing from the background (i.e., ancestral) regime (wAICc = 0.498), or *Phyllobates* and *H. infraguttatus* each having regimes separate from the background (wAICc = 0.241). None of the remaining models carried AICc weights above 0.1 (Fig. 4, Table 1).

**Figure 4.**
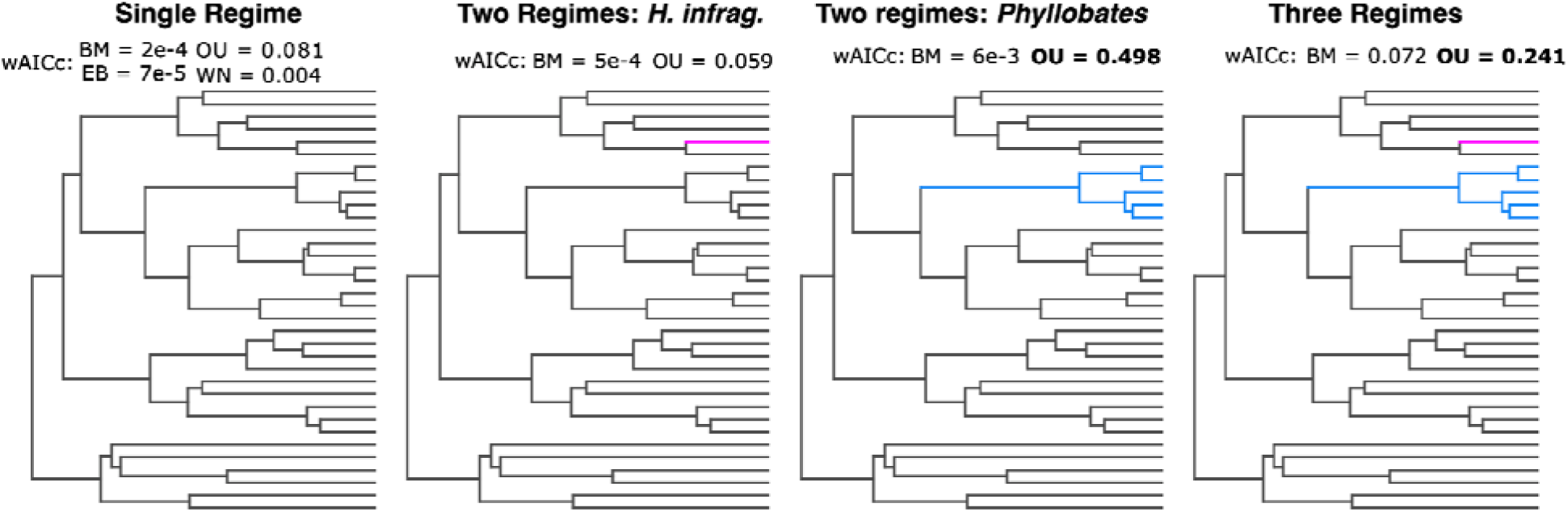
Models of continuous trait evolution tested for genome size in dendrobatids, along with their AICc weights. Branches evolving under the background evolutionary regime are colored gray, and subtrees evolving under distinct regimes are colored blue (*Phyllobates*) or magenta (*H. infraguttatus* and *H. elachyhistus*). The AICc for each regime configuration under different model types (i.e., BM, OU, etc.) are presented above each tree, and the two best-fitting models are highlighted in bold. All ten models were compared simultaneously. wAICc: Corrected Akaike weight; BM: Brownian Motion; EB: Early Burst; OU: Ornstein–Uhlenbeck; WN: White Noise.

### Correlates of genome size

Six of the 15 tested traits were correlated with genome size (Table 2). Body size, oocyte number, clutch size, and cell area showed positive correlations with genome size. Active metabolic rate (AMR; all metabolic variables measured in VO_2_ ml*h^-1^) and aerobic scope were negatively correlated with genome size, after controlling for body mass and phylogeny, while basal metabolic rate (BMR) was not. Because aerobic scope measures the difference between active and basal metabolic rates, this suggests that the correlation between scope and genome size is driven mainly by the active metabolic rate. Excluding *Phyllobates* from the models resulted in lower effect sizes and weaker statistical significance, as expected by the removal of 2-5 data points near the edges of trait distributions. Yet, the magnitude and direction of the correlations were qualitatively similar, indicating that the obtained correlations are not entirely driven by species in this genus. Regression parameters and associated statistics for the 15 predictors, including and excluding *Phyllobates*, are given in Table 2, and Figure 5 shows the relationships between genome size and the six correlated predictors.

**Figure 5.**
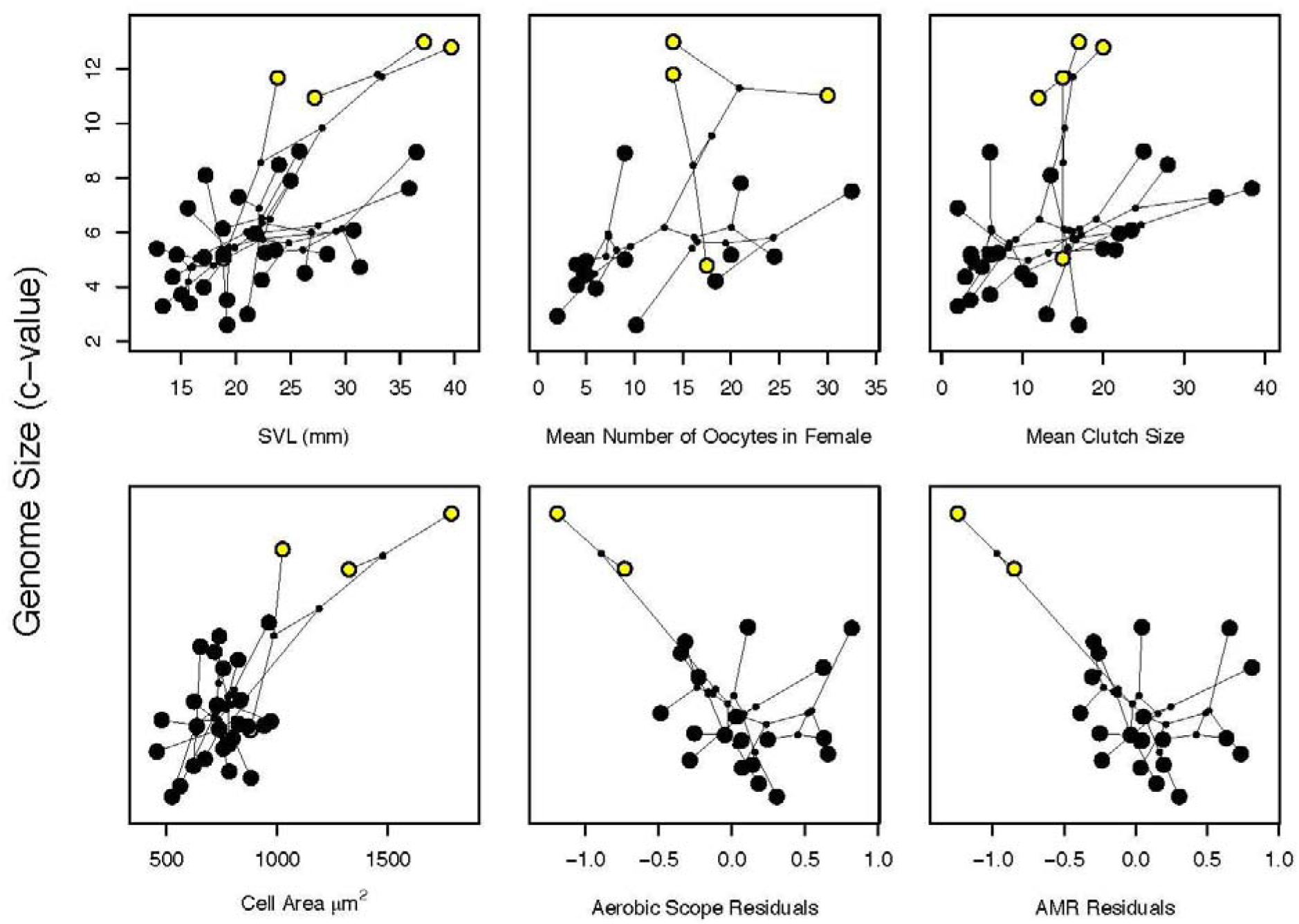
Phylomorphospace plots of the relationships between genome size and the six predictors that showed significant correlations. Large dots represent extant species, connected by lines that depict phylogeny. Small dots indicate bivariate ancestral state reconstructions. Species in the genus *Phyllobates* are colored yellow and all others black. For the two metabolic variables (aerobic scope and AMR) we plot the linear residuals with mass instead of the raw values to provide a visualization that controls for body mass. For model-fitting, mass was used as a covariate instead.

## Discussion

### Genome size evolution in Dendrobatidae

Dendrobatid genome sizes vary across the full range characterized in Anura, with a minimum C-value of 0.8 pg (*Aromobates tokuko*) and a maximum value of 13.0 pg (*Phyllobates bicolor*). The greater than 10-fold variation in genome size among dendrobatids (with N = 35 species) is broader than what has been observed in the few other similarly well-sampled amphibian families, such as plethodontid salamanders with a 5.2-fold range (13.90 – 71.65 pg; N = 95 species), hylid tree frogs with a 4.3-fold range (1.54 – 6.61 pg; N = 41 species), or bufonid toads with a 2.5-fold range (2.91 – 7.31 pg; N = 31 species) (Gregory 2022). Given the lack of comprehensive phylogenetic sampling in these and other families, we expect that additional cases of broad variation in genome size at lower taxonomic scales among amphibians will be described in the future.

The apparent occupation by dendrobatids of the entire range of genome sizes covered by frogs in just under 40 million years could be an artifact of the general lack of data for anuran genome sizes, as noted above. Alternatively, it may be explained by their much faster rate of genome size evolution (Fig. 2B), compared to the average rates across all frogs and other amphibian orders, which are over one order of magnitude lower (Liedtke et al. 2018). Previous work suggests that, across amphibians, genome size has evolved in a mostly gradual manner, with rare but pronounced saltational genome size shifts (Liedtke et al. 2018). Our results, which focused on a shallower phylogenetic scale, point to a somewhat similar scenario, with key differences. Phylogenetic comparative analyses provided good support for one or two discrete regime shifts in the evolution of genome size, although single-regime models were also supported in some cases (i.e., BAMM, *l1ou* under the pBIC), raising the possibility that the signature for discrete shifts come from clades at the extremes of the distribution of a quickly-evolving trait. Considering the number of species analyzed is on the low end for some of these analyses, we refrain from drawing definitive conclusions in this regard.

Irrespective of the number of independent evolutionary regimes, OU models were unambiguously supported over all others tested, suggesting that the diversification of genome sizes across Dendrobatidae has likely been marked by the existence of relatively stable “attractors”, such as adaptive optima, towards which lineages are drawn along their evolutionary trajectories. Under this framework, a regime shift constitutes a shift in the adaptive optimum, which results in evolution towards the new optimum due to positive selection (i.e., adaptation), followed by stabilizing selection around it. This contrasts with the patterns obtained across amphibian species, where lineages were found to gradually explore phenotypic space in a Brownian fashion, occasionally experience rapid jumps in genome size, and then continue their Brownian walk, suggesting that within Dendrobatidae genome size may have experienced stronger stabilizing selection than the general trend across Amphibia. Whether this is due to dataset characteristics, such as the temporal scales involved (∼40 vs. 350 MY), the degree of phylogenetic sampling (34 species within a family vs. 464 species across three orders) or the range of genome sizes investigated (0.8–13.0 pg vs. 0.8–121.0 pg), or it actually reflects dendrobatids having distinct evolutionary dynamics from the majority of other amphibian lineages remains to be clarified. Phylogenetic work with finer taxon sampling, as well as mechanistic studies on the effects of genome size change in different taxa would be especially insightful.

### Phenotypic correlates of genome size

We identified a significant correlation between genome size and six fitness-related traits: active metabolic rate (AMR), metabolic scope, snout-vent length (SVL), cell size, number of oocytes, and clutch size. Correlations with metabolic scope and the number of oocytes are not significant when we exclude the 5 *Phyllobates* lineages, yet qualitatively similar trends persist, and p-value changes are small (metabolic scope: p = 0.0496 vs. 0.054; number of oocytes: 0.0498 vs. 0.071). These changes are likely a combination of lower power (e.g., 18 vs 14 samples for oocyte number) and the removal of some of the extreme values that *Phyllobates* species represent in the dataset. In any case, similar traits to those in question, namely AMR and clutch size, remained correlated to genome size regardless of the species involved, so our conclusions are robust to these small statistical variations.

Amongst these correlates, all but AMR and scope were positively correlated with genome size. AMR and scope are thought to increase with origins of chemical defense (Santos and Cannatella 2011), while body size (SVL) may be of particular significance in poison frogs because it positively correlates with toxicity, bright coloration, and mating call complexity (Saporito et al. 2010; Santos and Cannatella 2011; Santos et al. 2014; Márquez et al. 2020). Despite our finding that three traits of the aposematic syndrome correlate with genome size, we found no relationship between alkaloid sequestration itself and genome size. AMR, scope, and SVL were correlated with genome size across our entire dataset of toxic and non-toxic species, even when toxicity is included as a covariate, further suggesting that the statistical relationship between these traits and genome size is unrelated to chemical defense.

The mechanistic link between genome size and metabolic rate is unclear, although it has been suggested to be mediated through cell size (Cavalier-Smith 1982; Gregory 2001), which we also found to be positively correlated with genome size. Larger cells divide less frequently and have a lower surface-area-to-volume ratio and thus a slower rate of nutrient and gas exchange, both of which should decrease metabolic rate (Gregory 2001). However, this pattern is not consistent across taxa. A negative correlation between genome size and basal metabolic rate (BMR) has been observed multiple times in birds and in mammals, although with weak support (Vinogradov 1995; Gregory 2002; Kozłowski et al. 2003). A negative correlation between genome size and BMR has been observed across all of Lissamphibia, but not when Anura and Caudata are considered independently (Gregory 2003). In addition, AMR and genome size are negatively correlated in some salamanders, but only at stress-inducing temperatures (Licht and Lowcock 1991), and ploidy level and AMR are negatively correlated in tadpoles but not froglets of the hybridogenetic frog *Pelophylax esculentus* (Hermaniuk et al. 2017). Finally, genome size does not appear to be associated with evolutionary saltations in BMR across extant vertebrates (Gardner et al. 2020), although there are exceptions in certain mammalian lineages (Uyeda et al. 2017). While AMR was not considered in some of these studies, potentially due to lack of data availability, our observation that AMR and metabolic scope are negatively correlated with genome size (controlling for mass) is fairly novel amongst amphibians. Dendrobatids exhibit highly specialized life history and physiology, including significant investment in parental care, coloration, toxin sequestration, and diet, all of which could potentially impact variation in metabolic rate. Additionally, AMR (but not BMR) is positively correlated with molecular evolution rate in Dendrobatidae (Santos 2012). Thus, studies of Dendrobatidae may present an opportunity to clarify the mechanistic link between genome size and metabolic rate.

Similarly to metabolic rate, the positive correlation between body size and genome size is thought to be mediated through cell size, and, additionally, through cell number (Arendt 2007; Hessen et al. 2013). While the mechanism underlying the positive correlation between genome size and cell size remains unclear, it may be driven by variation in transcriptional output or stoichiometric relationships between DNA amount and cytoplasmic protein levels (Mueller 2015). Because cell size is positively correlated with SVL and genome size in our data set, selection on cell size or related attributes may mediate the positive correlation between body size and genome size in poison frogs. In smaller species that have relatively large genomes for their size compared to other vertebrates (e.g., *Ranitomeya*: 4.4 – 6.8 pg: Stuckert et al. 2021), cell size could limit cell number and constrain morphological complexity (i.e., biological size), as has been seen in other amphibians (Levy and Heald 2015; Womack et al. 2019; Decena-Segarra et al. 2020). One potential consequence of miniaturization when genome size is not concomitantly reduced is the reduced complexity of the visual nervous system and hindering of vision-based behaviors such as active foraging, as has been observed in salamanders that have very large genomes (>30 pg; Matsushima et al. 1989). Given that some dendrobatids are diet specialists that actively forage for ants and mites, and that this behavior is associated with chemical defense, the loss of active foraging could be of particular consequence in Dendrobatidae (Toft 1995; Santos and Cannatella 2011; Tarvin et al. 2024). Furthermore, visual communication is thought to play an important role in several aspects of poison frog biology (e.g., Summers et al. 1999; Pröhl 2005; Yang et al. 2019). Thus, natural selection for the maintenance of body structures used in vision could play a role in genome size reduction in some dendrobatid species. We note, however, that the genera with the smallest genomes in our dataset, *Hyloxalus* and *Aromobates*, are neither miniaturized, brightly colored, nor outliers with respect to AMR (a correlate of active foraging; Santos and Cannatella 2011), thus we do not find strong support for this dynamic amongst the lineages we sampled.

Finally, we note that differences in genome size among poison frog species may be associated with differences in the activity of TEs, which appear to be currently proliferating in the genomes of at least two species that we examined (Rogers et al. 2018; Dittrich et al. 2024). Large genomes are thought to be hazardous in terms of fitness because of their larger mutation target, slower replication rates, and impacts on biological size. However, our understanding of the dynamics of genome size evolution are built on a small portion of living species, and it is plausible that amphibians, given their large genome sizes, may operate under different rules. For example, genome size is expected to increase in small populations where drift has a greater impact than selection on the fate of genetic variation. However, range size, a correlate of population size, is unrelated to genome size in salamanders (Rios-Carlos et al. 2024). In some salamanders, genome size is unrelated to TE deletion rate or genome-wide selection efficiency, which are both expected to decrease with larger genomes under current theory (Rios-Carlos et al. 2024; Wang et al. 2024). Differences in recombination rates (Rios-Carlos et al. 2024), DNA loss rates (Sun and Mueller 2014; Wang et al. 2024), and TE activity may help further explain why so many amphibians, including some poison frogs, have large genomes.

### Limitations

The wealth of formalin-fixed specimens in natural history collections around the world represent one of the richest and most complete resources to study phenotypic diversity — including genome size — in natural populations. However, chemicals used during specimen preservation such as formaldehyde and ethanol can affect nuclei compaction and Schiff’s reagent stain intensity. Museum specimens are usually fixed using a 4% solution of formaldehyde then stored long-term in 70% ethanol. Although formalin- and alcohol-fixation is also part of the Feulgen staining process, an empirical comparison of genome size estimates using Feulgen staining of fixed versus fresh tissues would help establish possible impacts of fixation and storage. Until these impacts are known, we recommend that, when precise estimates are required, genome size estimates from formalin-fixed tissues should be confirmed using fresh tissues and/or additional biological replicates. Despite potential caveats of using formalin-fixed tissues, we believe that any introduced errors are unlikely to overturn the general evolutionary patterns we identified in this study.

## Conclusion

The drivers of genome size evolution in eukaryotes remain a field of active research in evolutionary biology. Our results show that in dendrobatid poison frogs, genome size has evolved at much faster rates than in other amphibians — to the point of encompassing the entire range of genome sizes observed across anurans to date. Phylogenetic analyses suggest that this rapid evolution has proceeded primarily in a gradual manner and at a constant rate, constrained by purifying selection, and possibly with rare but appreciable events of punctuated evolution, although evidence for the latter is not definitive. Correlations between genome size and life history traits suggest that cell size may mediate changes in genome size within the group. Mechanisms underlying the relationship between cell size, metabolic rate, and genome size are not well understood, yet wide variation in each of these traits in Dendrobatidae could make the group suitable to provide mechanistic insight with future studies. Together these patterns suggest roles both for adaptive and neutral processes in shaping genome size evolution in Dendrobatidae. Our study contributes to understanding the multidimensional drivers of genome size, and sets the stage for future investigation into mechanisms driving genome size evolution in vertebrates.

## Data Accessibility

Raw data files, R scripts and detailed metadata are available for download from the Dryad Digital Repository http://doi.org/10.5061/dryad.0cfxpnw7d.

## Table Legends

**Table 1. Phylogenetically-aware linear regression coefficients of genome size against 15 morphological, metabolic, and life history traits, including or excluding species in the genus *Phyllobates*. Body mass was included as a covariate in models involving metabolic rates.**

**Table 2. Fit of various models of continuous trait evolution to genome size data in Dendrobatidae. Models are ranked based on AICc weight.**

## Footnotes

1 We could not find an R^2^ value reported in this study, but the authors report estimating genome size within 10% or 20% of published values for most of their standard species.

