## Supplementary Materials for "Genome size evolution and phenotypic correlates in the poison frog family Dendrobatidae"

**Supplementary Methods**

*Specimen identification*

We first reviewed the species identification for the museum specimens used in this study (Table S1) by examining the morphology and collection site and comparing them to relevant literature and occurrence databases. We generated a list of dendrobatid species that occur in the country of a given specimen’s collection site using the AmphibiaWeb country-specific species search (AmphibiaWeb 2024). Dendrobatid morphology is difficult to confuse with other co-occurring species, but when relevant we considered non-dendrobatids (e.g., *Lithodytes lineatus, Pristimantis* spp.) as potential misidentifications. We excluded from the country-specific species list any dendrobatid whose range is unlikely to overlap with the specimen collection site based on relevant literature and publicly available collection data (e.g., GBIF data imported into AmphibiaWeb’s Berkeley Mapper tool). Because range data present in online databases is imperfect, when necessary we also reviewed primary literature to validate specific identifications. We excluded any co-occurring species that could be clearly ruled out with morphological characters (according to Grant et al. 2006, 2017) and then reviewed each specimen using the dendrobatid character matrix from Grant et al. (2017) to either confirm or revise each putative ID (see Table S1 for more details).

We also reviewed the source literature for all of the collated genome size measurements of anurans in the Animal Genome Size Database (AGSD; Gregory 2022). For non-dendrobatid anurans on AGSD, we minimally checked that the species identification listed matched with the collection site of the specimen (when data were available). For dendrobatid data published on AGSD, when possible we additionally reviewed source specimens or photographs of source specimens to confirm species identifications (see Table S2).

*Flow Cytometry*

Given the variation that sex, strain and storage conditions can introduce into commercially available chicken red blood cells (CRBCs), and that flow cytometry estimates of genome size are more accurate when the standard peak falls near but not on the sample peak (Hare and Johnston 2011), we opted to use insect muscle tissue as a standard that has been scored against freshly collected CRBCs (Bennett et al. 2003). Nuclei were released from two species that serve as standards for genome size estimates (*Drosophila virilis* [1C = 328] and *Periplaneta americana* [1C = 3303]) by combining a *D. virilis* female head and part of a *P. americana* male head in 1 ml of cold Galbraith buffer in a 2-ml Dounce and grinding with 10 gentle strokes using an A (loose) pestle. Nuclei were next released from a 1-cm^3^ piece of *Phyllobates* leg or tadpole tail muscle in 25 ul of cold Galbraith buffer using 25 chops of tissue with a fresh single edged razor blade (one species at a time). The *Phyllobates* nuclei were diluted to 0.95 ml with additional cold Galbraith buffer and filtered through 45 um nylon mesh into a 1.5 mL microfuge tube along with 0.05 ml of the cold Galbraith buffer pulled from the Dounce with the nuclei from the standards. The combined nuclei from the chopped poison frog muscle and the ground *D. virilis* *and P. americana* standards were stained with 25 µl of propidium iodide (1 mg/ml) for 3 hours in the dark at 4 ℃, after which time the mean red PI fluorescence of the stained DNA in the nuclei of the sample and standards was quantified using a CytoFlex flow cytometer (Beckman Coulter). Haploid (1C) DNA quantity was calculated as (2C sample mean fluorescence/2C standard mean fluorescence) times 328 Mbp for the *D. virilis* standard and times 3303 Mbp for the *P. americana*. The two standards produced very similar average genome size estimates, although the standard error was larger for the *D. virilis* standard. Thus, the reported 1C genome size estimate for *Phyllobates* species is based on the estimate using the *P. americana* standard. We quantified genome size for 2-5 individuals per species and used the average of individual genome sizes in further analyses. A single male *P. bicolor* individual was found to have roughly twice the genome size as its conspecifics. This frog had noticeable deformities in some of its bones, and was unable to successfully fertilize egg clutches, which leads us to suspect it may have had an anomalous ploidy, and exclude its genome size estimate from average calculations. The raw genome size data for all individuals are provided in the Dryad repository (Table_S2_exp.tsv in the Datasets folder).

**Supplementary Results**

*Specimen Identification and Reassignment*

We reviewed the identification of 66 specimens from the MVZ representing 38 species, including 27 species of Dendrobatidae (Table S1). Excluding six specimens whose identifications were not yet updated to new taxonomy (e.g., *Limnodynastes ornatus* is now *Platypectrum ornatum*, *Bufo cognatus* is now *Anaxyrus cognatus*, and *Elachistocleis ovalis* is a *nomen dubium* (see notes in Heyer and and Liem 1976; Caramaschi 2010; Barrio-Amorós et al. 2019)), 18 (27%) of the specimens were identified incorrectly to genus, species, or both (Table S1). Many species of dendrobatid frogs are morphologically similar including some mimics and many inconspicuously colored species, which makes correct identification difficult, especially after long-term storage in ethanol. Thus, we note some remaining uncertainty regarding specific specimens (see details in Table S1).

We reviewed 48 genome size estimations of 19 species published on the Animal Genome Size Database (Table S2), including 8 measurements of 8 dendrobatid species. Five total records (10%) were identified incorrectly to genus, species, or both (4 were dendrobatids). One (*Dendrobates femoralis*) had since been updated in VertNet as *Lithodytes lineatus* (a frog from a different family) but had not yet been updated in AGSD. With specimen photographs we were able to reassign two other samples (*Epipedobates trivittatus, Colostethus marchesianus*) to their correct identifications (*Allobates femoralis, Anomaloglossus* cf. *surinamensis,* respectively), which were both phylogenetically distant from the original identifications.

Specimens examined in MacCulloch et al. (1996) were assigned at the time to two morphotypes of *Epipedobates pictus* (*E.* cf. *pictus* sp. 1 and *E.* cf. *pictus* sp. 2), and since reidentified as *Ameerega hahneli* (Reynolds and MacCulloch 2012). When we were reviewing these specimens, two phenotypic groupings based on light and dark coloration corresponded to the sample sizes described by MacCulloch et al. (1996); thus, we assume that *E.* cf. *pictus* sp. 1 corresponds to lighter-colored specimens with museum numbers ROM 22797-22799 and that *E.* cf. *pictus* sp. 2, corresponds to darker-colored specimens with museum numbers ROM 22795-22796 and ROM 22805-22807. In contrast to the re-identification by Reynolds and MacCulloch (2012), we observed that *E.* cf. *pictus* sp. 2 individuals have a body shape more similar to *Allobates femoralis* (uniform trunk width) than *Ameerega hahneli* (pear-shaped), which is further corroborated by morphological characteristics described by Grant et al. (2017): the *E.* cf. *pictus* sp. 2 specimens possessed Finger V length (character 5) as state 1 (reaching distal half of distal subarticular tubercle of Finger IV), yet *Ameerega hahneli* is expected to be state 0 (Finger V length surpassing distal subarticular tubercle of Finger IV). The animals assigned to *E.* cf. *pictus* sp. 1 were small and might be juveniles of the same or another species (but likely not *Ameerega hahneli*, assuming that character 5 state doesn’t change with ontogeny). Based on our review of the specimens, we tentatively reassign *E.* cf. *pictus* sp. 2 to *Allobates femoralis* but refrain from assigning a species identification for *E.* cf. *pictus* sp. 1 without further information and we do not include them in our analyses.

*Potential genome size polymorphism or mimicry in two dendrobatid lineages*

Genome size estimates and specimen identifications suggest either genome size polymorphism or extreme mimicry between *Ameerega hahneli* and *Allobates femoralis*, or cryptic species of *Am. hahneli*. Specifically, the genome size of two specimens from Colombia (MVZ:Herp:63120 and MVZ:Herp:63759) that were collected at the same site and were morphologically indistinguishable from each other and from additional specimens collected at a nearby locality, which were identified as *Am. hahneli* (UQUINDÍO:HERPETOS-UQ:ARUQ-1099–1102; Table S1) based on mitochondrial DNA sequences, were notably different: 5.24 pg and 7.80 pg, respectively (Table S5). Similarly, MacCulloch et al. (1996) also reported two C-values (6.55 pg and 8.9 pg) for specimens collected in Guyana that were initially identified as *Epipedobates* cf. *pictus* sp. 1 and *E.* cf. *pictus* sp. 2*.* We reassigned the latter to *Allobates femoralis*, but we could not unequivocally identify the former (see Table S2 for details). Camper et al. (1993) reported the C-value of an *Allobates femoralis* specimen collected from French Guiana (previously identified as *Epipedobates trivittatus*) as 8.49 pg, similar to the value from MacCulloch et al. (1996) for *E.* cf. *pictus* sp. 2, and supporting our reassignment of *E.* cf. *pictus* sp. 2 to *Al. femoralis*. However, such disparate values from two potential conspecifics collected at the same locality is perplexing. Three scenarios are plausible. 1) MVZ:Herp:63759 is misidentified and is actually *Al. femoralis*, suggesting that *Al. femoralis* varies in genome size (with C-values of 7.8 pg, 8.49 pg, 8.9 pg). 2) Both MVZ specimens are identified correctly as *Am. hahneli*, suggesting polymorphism in *Am. hahneli* genome size (with C-values of 5.24 pg and 7.8 pg). 3) The two MVZ specimens represent cryptic species with distinct genome sizes. Unfortunately, the MVZ specimens were collected in 1950 and well-preserved tissue samples are unavailable to easily verify these patterns.

*Accuracy of taxonomic identification in public databases*

A meaningful proportion of data in each of the databases we used (10% in AGSD, 27% in MVZ) was found to be associated with an incorrect or putatively incorrect species identification (all identifications for specimens included in this paper will be updated in AGSD and MVZ during the review process). We note, however, that most of our taxonomic reidentifications pertained to data from 2 (Camper et al. 1993; MacCulloch et al. 1996) of 25 reviewed papers and therefore frequency of taxonomic inaccuracies with encountered may not represent the accuracy of the AGSD as a whole. We caution and encourage users of public data to take the time to validate species identifications. Validation may be especially necessary when organisms were collected by scientists that are not experts in the focal taxon, when the specimen identifications have not been revised for several decades, and for taxa with large amounts of morphologically similar taxa. All three of these criteria apply to the MVZ dendrobatid specimens. The prevalence of misidentified specimens in databases is not new or surprising (Szczęsny and Godunko 2007; Mulcahy et al. 2022). DNA barcoding is one approach that can validate museum specimen identifications (Hebert et al. 2003), though it may not always be accurate if genetic structure at the used marker(s) does not correlate perfectly with species identity (Chambers and Hebert 2016; Mulcahy et al. 2022) or database content. Digitizing collections can also accelerate the process of validating identifications by increasing access to specimens (Cook et al. 2014).

**Supplementary Figures**

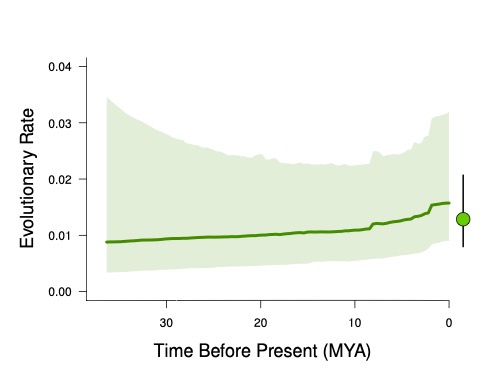

**Figure S1.** Comparison of tree-wide evolutionary rate over time estimates from BAMM, and the tree-wide rate estimated under a single-regime Brownian Motion model fit using *geiger*. The line and surrounding polygon represent the median and 95% highest credibility interval from the BAMM post-burnin posterior distribution. The point and bar correspond to the point estimate and 95% confidence interval from the *geiger* maximum-likelihood optimization. The confidence interval was obtained from the model’s Hessian matrix. Considering the BAMM estimate at any given timeslice is across the entire tree, and that the evidence for distinct evolutionary regimes is restricted to a few branches, a single-rate model should most closely approximate BAMM’s mixed model.

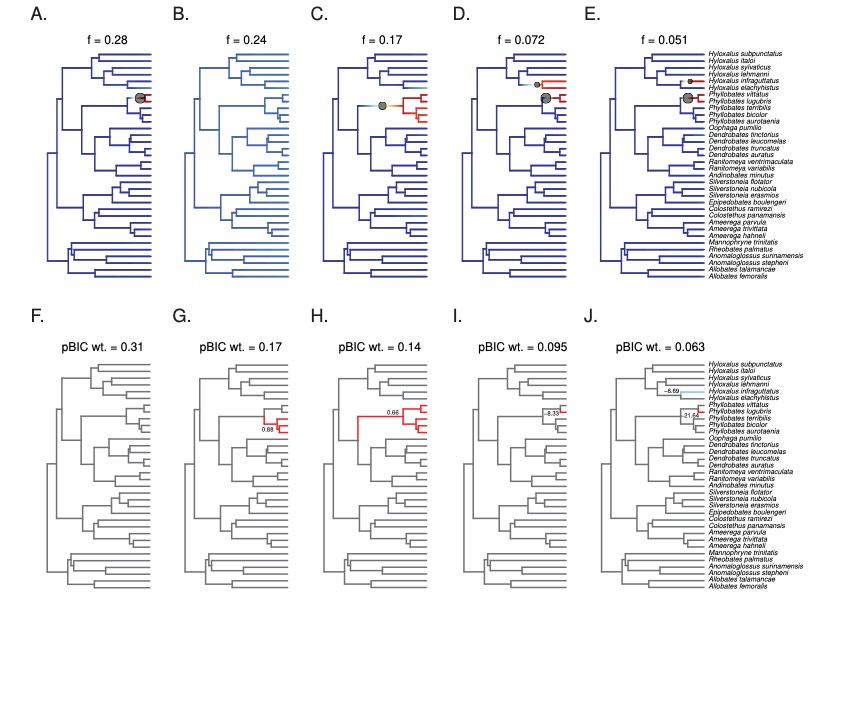

**Figure S2.** Evidence for discrete shifts in genome size evolution. Panels A-E show the five most frequent rate shift configurations in the BAMM posterior output along with their posterior probabilities. Circles mark positions of rate shifts, with sizes proportional to marginal shift probability. Warmer colors represent faster evolutionary rates. These configurations account for 81.3% of the posterior density. Panels F-J show the five best-fitting multi-regime OU models from *l1ou* and their pBIC weights. Clades are colored by OU regime, and branch numbers indicate the magnitude of each shift in the subtree’s “optimal” phenotype. These models correspond to 78% of the pBIC weight among all tested models.

**Table S1**. Specimens used for genome size estimation, with notes on revised species identifications and other specimen-specific information.

| Species | batch | Museum Number | Latitude | Longitude | Locality, verbatim | Coord. error (km) | Elev. (m) | Elev. Source | Effect of error on elevation estimate | Collection date | SVL (mm) | Age | Original species ID | Validation of ID |
| --- | --- | --- | --- | --- | --- | --- | --- | --- | --- | --- | --- | --- | --- | --- |
| *Allobates femoralis* | 2 | MVZ:Herp:172012 | -5.61667 | -78.65 | 2 mi inland from Bella Vista on Caijaru River, South America, Peru, Loreto | 7.11 | NA | NA | NA | 9 Sep 1974 | 23.93 | adult | *Allobates femoralis* | MVZ:Herp:172012 was originally identified as *Allobates femoralis*. The only other morphologically similar sympatric species is *Ameerega hahneli*. Toe pad and finger morphology support *A. femoralis* over *A. hahneli*. We maintained original ID as *A. femoralis* |
| *Allobates talamancae* | 2 | MVZ:Herp:113731 | 8.69745 | -83.48731 | vicinity of Tropical Science Center Field Station, Rincon de Osa, Central America, Costa Rica, Puntarenas | 1.20 | NA | NA | NA | 3 Jul 1974 | 20.1 | adult | *Allobates talamancae* | MVZ:Herp:113731 was originally identified as *Allobates talamancae*. The only other likely sympatric species is *Silverstoneia nubicola*. Presence of dorsolateral stripe, dark ventral coloration, and finger/toe morphology support *A. talamancae* over *S. nubicola*. We maintained the original ID as *Allobates talamancae* |
| *Allobates talamancae* | 2 | MVZ:Herp:113732 | 8.69745 | -83.48731 | vicinity of Tropical Science Center Field Station, Rincon de Osa, Central America, Costa Rica, Puntarenas | 1.20 | NA | NA | NA | 3 Jul 1974 | 20.35 | adult | *Allobates talamancae* | MVZ:Herp:113732 was originally identified as *Allobates talamancae*. The only other likely sympatric species is *Silverstoneia nubicola*. Presence of dorsolateral stripe, dark ventral coloration, and finger/toe morphology support *A. talamancae* over *S. nubicola*. We maintained the original ID as *Allobates talamancae* |
| *Ameerega hahneli* | 1, 2 | MVZ:Herp:63120 | 3.5 | -73.833333 | 46 km S and 22 km W San Martin, South America, Colombia, Meta | 2.00 | 488 | MVZ | small, 100m range in elevation in region | 25 Nov 1950 | 19.9 | adult | *Epipedobates pictus* | MVZ:Herp:63120 was originally identified as Epipedobates pictus (now Ameerega). Specimen is morphologically similar to Ameerega hahneli specimens collected nearby in Caquetá, Colombia currently in the Universidad de Quindío herp collection (UQUINDÍO:HERPETOS-UQ:ARUQ-1099, UQUINDÍO:HERPETOS-UQ:ARUQ-1100, UQUINDÍO:HERPETOS-UQ:ARUQ-1101, UQUINDÍO:HERPETOS-UQ:ARUQ-1102, whose identification as Ameerega hahneli was confirmed by COI sequencing). Identification was revised to Ameerega hahneli. |
| *Ameerega hahneli* | 1 | MVZ:Herp:63759 | 3.5 | -73.833333 | 46 km S and 22 km W San Martin, South America, Colombia, Meta | 2.00 | 488 | MVZ | small, 100m range in elevation in region | 25 Nov 1950 | 23.63 | adult | *Epipedobates pictus* | MVZ:Herp:63759 was originally identified as Epipedobates pictus (now Ameerega). Specimen is morphologically similar to Ameerega hahneli specimens collected nearby in Caquetá, Colombia currently in the Universidad de Quindío herp collection (UQUINDÍO:HERPETOS-UQ:ARUQ-1099, UQUINDÍO:HERPETOS-UQ:ARUQ-1100, UQUINDÍO:HERPETOS-UQ:ARUQ-1101, UQUINDÍO:HERPETOS-UQ:ARUQ-1102, whose identification as Ameerega hahneli was confirmed by COI sequencing). Identificiation was revised to Ameerega hahneli. |
| *Ameerega parvula* | 1 | MVZ:Herp:162946 | -4.45563 | -78.16123 | vicinity of Huampami (Aguaruna village), Rio Cenepa South America, Peru, Amazonas | 0.41 | 213 | MVZ | none | 9 Jul 1978 | 28.33 | adult | *Ameerega parvula* | MVZ:Herp:162946 was originally identified as *Ameerega parvula*. *A. bilinguis* and *A. zaparo* are both morphologically similar sympatric species. Relatively large body size and toe webbing support *A. parvula* over *A. bilinguis* and *A. zaparo*. We maintained the original ID as *A. parvula*. |
| *Ameerega parvula* | 1 | MVZ:Herp:162950 | -4.45563 | -78.16123 | vicinity of Huampami (Aguaruna village), Rio Cenepa South America, Peru, Amazonas | 0.41 | 213 | MVZ | none | 14 Jul 1978 | 24.26 | adult | *Ameerega parvula* | MVZ:Herp:162950 was originally identified as *Ameerega parvula*. *A. bilinguis* and *A. zaparo* are both morphologically similar sympatric species. Relatively large body size and toe webbing support *A. parvula* over *A. bilinguis* and *A. zaparo*. We maintained the original ID as *A. parvula*. |
| *Ameerega trivittata* | 1 | MVZ:Herp:247538 | 4.949833 | -55.186333 | Mazaroni Rd., 0.3 km from junction with Brownsberg Rd., Brownsberg Nature Park, South America, Suriname, Brokopondo District | 0.30 | 495 | MVZ | none | 26 Mar 2004 | 36.45 | adult | *Epipedobates pictus* | MVZ:Herp:247538 was originally identified as *Epipedobates pictus* (now *Ameerega*). Specimen body size is significantly larger than the expected size range of A*. picta* as well as other sympatric dendrobatid species. Large specimen size and coloration matches expectation for *Ameerega trivittata.* |
| *Ameerega trivittata* | 1 | MVZ:Herp:247537 | 4.9505 | -55.183333 | Brownsberg Nature Park Headquarters, South America, Suriname, Brokopondo District | 0.30 | 495 | MVZ | none | 20 Mar 2004 | 35.2 | adult | *Epipedobates pictus* | MVZ:Herp:247537 was originally identified as *Epipedobates pictus* (now *Ameerega*). Specimen body size is significantly larger than the expected size range of A*. picta* as well as other sympatric dendrobatid species. Large specimen size and coloration matches expectation for *Ameerega trivittata.* |
| *Anaxyrus cognatus* | 2 | MVZ:Herp:65609 | 35.208 | -101.46937 | 5 mi W Conway on Rte. 66, North America, United States, Texas, Carson County | 7.60 | NA | NA | NA | 8 Aug 1954 | 23.99 | adult | *Bufo cognatus* | MVZ:Herp:65609 was originally identified as *Anaxyrus cognatus.* The specimen morphologically and collection site matches that of *A. cognatus*. We mainained the original identification as *Anaxyrus cognatus*. |
| *Andinobates minutus* | 2 | MVZ:Herp:210477 | 9.26667 | -78.96667 | Nusagundi, Kuna Yala, Central America, Panama | 11.10 | 250 | MVZ | moderate, elevation range in area 200-400 | 19 Oct 1988 | 13.32 | adult | *Andinobates minutus* | MVZ:Herp:210477 was originally identified as *Adinobates minutus*. *A. fulguritis* and *A. geminisae* are both symaptric with and morphologically similar to *A. minutus*. Absence of ventrolateral stripe and ventral patterning support *A. minutus* over *A. fulguritis* and *A. geminisae*. We maintained the original ID as *Andinobates minutus*. |
| *Anomaloglossus stepheni* | 2 | MVZ:Herp:247534 | 4.9505 | -55.183333 | Brownsberg Nature Park Headquarters, South America, Suriname, Brokopondo District | 0.03 | 495 | MVZ | variable in the region, but no error in GPS coord. | 27 Mar 2004 | 18.87 | adult | *Anomaloglossus stepheni* | MVZ:Herp:247534 was originally identified as *Anomaloglossus stepheni*. Presence of a median lingual process (reviewed via photos of the specimen) as well as coloration and range support identification as *Anomaloglossus stepheni, but there is some possibility that it is another member of the genus*. We maintained the original ID as *A. stepheni with some uncertainty*. |
| *Anomaloglossus stepheni* | 0 | MVZ:Herp:247535 | 4.9505 | -55.183333 | Brownsberg Nature Park Headquarters, South America, Suriname, Brokopondo District | 0.03 | 495 | MVZ | variable in the region, but no error in GPS coord. | 28 Mar 2004 | 19.51 | adult | *Anomaloglossus stepheni* | MVZ:Herp:247535 was originally identified as *Anomaloglossus stepheni*. Presence of a median lingual process (reviewed via photos of the specimen) as well as coloration and range support identification as *Anomaloglossus stepheni,* but there is some possibility that it is another member of the genus. We maintained the original ID as *A. stepheni* with some uncertainty. |
| *Bombina bombina* | 2 | MVZ:Herp:164719 | 47.85833 | 16.8761 | 3.2 km E Podersdorf, Europe, Austria, Burgenland | 6.91 | NA | NA | NA | 28 June 1978 | 44.7 | NA | *Bombina bombina* | MVZ:Herp:164719 was originally identified as *Bombina bombina.* The specimen morphologically and collection site matches that of *B. bombina*. We mainained the original identification as *Bombina bombina*. |
| *Bombina bombina* | 2 | MVZ:Herp:164720 | 47.85833 | 16.8761 | 3.2 km E Podersdorf, Europe, Austria, Burgenland | 6.91 | NA | NA | NA | 28 June 1978 | 40.54 | NA | *Bombina bombina* | MVZ:Herp:164720 was originally identified as *Bombina bombina.* The specimen morphologically and collection site matches that of *B. bombina*. We mainained the original identification as *Bombina bombina*. |
| *Bombina variegata* | 2 | MVZ:Herp:8380 | 46.074692 | 18.234205 | Pecs, Europe, Hungary, Baranya County | 10.00 | NA | NA | NA | 8 May 1904 | 38.71 | NA | *Bombina variegata* | MVZ:Herp:8380 was originally identified as *Bombina variegata.* The collection site matches that of *B. bombina* but the estimated genome size is distinct. We mainained the original identification as *Bombina variegata* with some uncertainty. |
| *Bombina variegata* | 2 | MVZ:Herp:8381 | 46.074692 | 18.234205 | Pecs, Europe, Hungary, Baranya County | 10.00 | NA | NA | NA | 8 May 1904 | 34.8 | NA | *Bombina variegata* | MVZ:Herp:8381 was originally identified as *Bombina variegata.* The collection site matches that of *B. bombina* but the estimated genome size is distinct. We mainained the original identification as *Bombina variegata* with some uncertainty. |
| *Colostethus panamansis* | 1 | MVZ:Herp:128605 | 8.61027 | -80.13976 | 1 km SW (by air) Hotel Greco, El Valle de Anton, Central America, Panama, Cocle | 6.20 | 630 | MVZ | moderate, range of elevation in region is 600-1000 | 27 Aug 1975 | 25.52 | adult | *Colostethus pratti* | MVZ:Herp:128605 was originally identified as *Colostethus pratti. Allobates talamancae, C. panamansis, C. pratti, C. latinasus, S. nubicola, Silverstoneia flotator a*re all morphologically similar sympatric species. An incomplete oblique lateral stripe on specimen of interest likely excludes *Allobates* and *Silverstoneia* genera as possible IDs. Altitudinal range likely excludes *C. latinasus.* Presence of preaxial webbing on toes I-IV supports *C. panamansis* over *C. pratti.* Presence of shoulder patch also supports ID as *C. panamansis* (Ibanez et al. 2017). ID revised to *Colestethus panamansis* with some uncertainty*.* |
| *Colostethus panamansis* | 1 | MVZ:Herp:128611 | 8.61027 | -80.13976 | 1 km SW (by air) Hotel Greco, El Valle de Anton, Central America, Panama, Cocle | 6.20 | 630 | MVZ | moderate, range of elevation in region is 600-1000 | 27 Aug 1975 | 26.04 | adult | *Colostethus pratti* | MVZ:Herp:12861 was originally identified as *Colostethus pratti. Allobates talamancae, C. panamansis, C. pratti, C. latinasus, S. nubicola, Silverstoneia flotator a*re all morphologically similar sympatric species. An incomplete oblique lateral stripe on specimen of interest likely excludes *Allobates* and *Silverstoneia* genera as possible IDs. Altitudinal range likely excludes *C. latinasus.* Presence of preaxial webbing on toes I-IV supports *C. panamansis* over *C. pratti.* Presence of shoulder patch also supports ID as *C. panamansis* (Ibanez et al. 2017). ID revised to *Colestethus panamansis* with some uncertainty*.* |
| *Dendrobates auratus* | 1 | MVZ:Herp:83069 | 8.48333 | -78.51667 | Isla Taboga Pacific Ocean, Gulf of Panama, Panama, Panama, Isla Taboga | 182.60 | 0 | RM_Approx | coordinates seem wrong; we used 0, as it is a low island. | 13 Oct 1967 | 24.54 | adult | *Dendrobates auratus* | MVZ:Herp:83069 was originally identified as *Dendrobates auratus*. *D. auratus* is the only known frog species on Isla Taboga, the collection site of this specimen. We maintained the originally identification as *Dendrobates auratus*. |
| *Dendrobates auratus* | 0 | MVZ:Herp:83077 | 8.4833 | -78.51667 | Isla Taboga Pacific Ocean, Gulf of Panama, Panama, Panama, Isla Taboga | 182.60 | 0 | RM_Approx | coordinates seem wrong; we used 0, as it is a low island. | 22 Oct 1967 | 20.82 | adult | *Dendrobates auratus* | MVZ:Herp:83069 was originally identified as *Dendrobates auratus*. *D. auratus* is the only known frog species on Isla Taboga, the collection site of this specimen. We maintained the originally identification as *Dendrobates auratus*. |
| *Dendrobates leucomelas* | 1 | MVZ:Herp:81475 | 6.33333 | -63.5 | Ucaima, South America, Venezuela, Bolivar | 403.60 | 427 | RM_Approx | Elevation varies between ~250-600 in the area. We used Ucayma at the foot of a tepui (D. leuco habitat) | 2 Feb 1967 | 33.38 | adult | *Dendrobates leucomelas* | MVZ:Herp:81475 was originally identified as *Dendrobates leucomelas*. Color pattern, size, and body shape all strongly support *D. leucomelas* identification. Other sympatric dendrobatids are not morphologically similar. We maintained the original identification as *Dendrobates leucomelas.* |
| *Dendrobates leucomelas* | 1 | MVZ:Herp:81474 | 6.33333 | -63.5 | Ucaima, South America, Venezuela, Bolivar | 403.60 | 427 | RM_Approx | Elevation varies between ~250-600 in the area. We used Ucayma at the foot of a tepui (D. leuco habitat) | 30 Jan 1967 | 29.3 | adult | *Dendrobates leucomelas* | MVZ:Herp:81474 was originally identified as *Dendrobates leucomelas*. Color pattern, size, and body shape all strongly support *D. leucomelas* identification. Other sympatric dendrobatids are not morphologically similar. We maintained the original identification as *Dendrobates leucomelas.* |
| *Dendrobates truncatus* | 1 | MVZ:Herp:42005 | 3.8 | -75.5 | 8 km N Chaparral, South America, Colombia, Tolima | 2.00 | 700 | MVZ | small, only 200m range in the 2-km radius | 1 Dec 1944 | 28.35 | adult | *Dendrobates truncatus* | MVZ:Herp:42005 was originally identified as *Dendrobates truncatus*. *D. truncates* is sympatric with species of *Andinobates*, *Oophaga*, and *Phyllobates*. However, the specimen is larger than the expectation for *Adinobates*, and its coloration is distinct from that of *Oophaga* and *Phyllobates*. We maintained the original identification as *D. truncates*. |
| *Elachistocleis pearsei* | 1, 2 | MVZ:Herp:210460 | 8.71667 | -79.9 | Cerro Campana, Central America, Panama, Panama | 4.10 | 720 | MVZ | potentially large, elevation ranges 200-800 in region | 7 Nov 1988 | 17.02 | NA | *Elachistocleis ovalis* | EXCLUDE. This specimen was originally identified as Elachistocleis ovalis, but this specific epithet is invalid. Based on the collection locality, this specimen may belong to *E. pearsei* (Jowers et al. 2021) |
| *Elachistocleis pearsei* | 2 | MVZ:Herp:210459 | 8.71667 | -79.9 | Cerro Campana, Central America, Panama, Panama | NA | NA | MVZ | potentially large, elevation ranges 200-800 in region | 7 Nov 1988 | 34.61 | NA | *Elachistocleis ovalis* | EXCLUDE. This specimen was originally identified as Elachistocleis ovalis, but this specific epithet is invalid. Based on the collection locality, this specimen may belong to *E. pearsei* (Jowers et al. 2021) |
| *Elachistocleis pearsei* | 0 | MVZ:Herp:210457 | 8.71667 | -79.9 | Cerro Campana, Central America, Panama, Panama | 4.10 | 720 | MVZ | potentially large, elevation ranges 200-800 in region | 26 Oct 1988 | NA | NA | *Elachistocleis ovalis* | EXCLUDE. This specimen was originally identified as Elachistocleis ovalis, but this specific epithet is invalid. Based on the collection locality, this specimen may belong to *E. pearsei* (Jowers et al. 2021) |
| *Epipedobates boulengeri* | 0 | MVZ:Herp:192363 | 3.7 | -77 | 6-8 km W Danubio on old road from Cali to Buenaventura, near La Chorrera, South America, Colombia, Valle del Cauca | 2.00 | 450 | MVZ | none | 21 Jul 1977 | 11.19 | adult | *Epipedobates boulengeri* | MVZ:Herp:192363 was originally identified as *Epipedobates boulengeri.* The specimen's mottled venter and general morphology rules out other potential co-occuring species. We mainained the original identification as *Epipedobates boulengeri*. |
| *Epipedobates boulengeri* | 1 | MVZ:Herp:192359 | 3.7 | -77 | 6-8 km W Danubio on old road from Cali to Buenaventura, near La Chorrera, South America, Colombia, Valle del Cauca | 2.00 | 450 | MVZ | none | 21 Jul 1977 | 12.79 | adult | *Epipedobates boulengeri* | MVZ:Herp:192359 was originally identified as *Epipedobates boulengeri.* The specimen's mottled venter and general morphology rules out other potential co-occuring species. We mainained the original identification as *Epipedobates boulengeri*. |
| *Hyla versicolor* | 1, 2 | MVZ:Herp:60480 | 41.7038 | -86.2211 | 1 mi E Notre Dame campus, South Bend, North America, United States, Indiana, Saint Joseph County | 3.13 | NA |  | none | 24 May 1951 | 40.15 | NA | *Hyla versicolor* | MVZ:Herp:60480 was originally identified as *Hyla versicolor.* The specimen morphologically and collection site matches that of *H. versicolor*. We mainained the original identification as *Hyla versicolor*. |
| *Hyla versicolor* | 2 | MVZ:Herp:60474 | 41.7038 | -86.2211 | 1 mi E Notre Dame campus, South Bend, North America, United States, Indiana, Saint Joseph County | 3.13 | NA |  | none | 24 May 1951 | 42.52 | NA | *Hyla versicolor* | MVZ:Herp:60474 was originally identified as *Hyla versicolor.* The specimen morphologically and collection site matches that of *H. versicolor*. We mainained the original identification as *Hyla versicolor*. |
| *Hyloxalus elachyhistus* | 1 | MVZ:Herp:112482 | -3.98191 | -79.217 | 2 km N Loja, South America, Ecuador, Loja | 4.86 | 2100 | MVZ | small | 23 Jul 1971 | 16.03 | adult | *Hyloxalus elachyhistus* | MVZ:Herp:112482 is a paratype of *Hyloxalus elachyhistus*. |
| *Hyloxalus elachyhistus* | 0 | MVZ:Herp:112480 | -3.98191 | -79.217 | 2 km N Loja, South America, Ecuador, Loja | 4.86 | 2100 | MVZ | small | 23 Jul 1971 | 18.44 | adult | *Hyloxalus elachyhistus* | MVZ:Herp:112480 was collected contemporaneously with MVZ:Herp:112482, which is a paratype of *Hyloxalus elachyhistus*. |
| *Hyloxalus infraguttatus* | 1 | MVZ:Herp:77177 | -2.167 | -79.04599 | 95 km E Guayaquil on road to Quito, South America, Ecuador, Guayas | 74.45 | 3000 | MVZ | potentially large, site is right at foothill | 4 Mar 1964 | 20.31 | adult | *Hyloxalus infraguttatus* | MVZ:Herp:77177 was originally identified as *Hyloxalus infraguttatus. H. infraguttatus* is sympatric with the morphologically similar *Epipedobates machalilla* and *E. tricolor*. The presence of white spotting on the throat and venter favor *H. infraguttatus* over either *Epipedobates* species. We maintained the original identification as *Hyloxalus infragutattus*. |
| *Hyloxalus infraguttatus* | 0 | MVZ:Herp:77181 | -2.167 | -79.04599 | 95 km E Guayaquil on road to Quito, South America, Ecuador, Guayas | 74.45 | 3000 | MVZ | potentially large, site is right at foothill | 4 Mar 1964 | 18.11 | adult | *Hyloxalus infraguttatus* | MVZ:Herp:77181 was originally identified as *Hyloxalus infraguttatus. H. infraguttatus* is sympatric with the morphologically similar *Epipedobates machalilla* and *E. tricolor*. The presence of white spotting on the throat and venter favor *H. infraguttatus* over either *Epipedobates* species. We maintained the original identification as *Hyloxalus infragutattus*. |
| *Hyloxalus italoi* | 1 | MVZ:Herp:162627 | -4.45563 | -78.16123 | vicinity of Huampami (Aguaruna village), Rio Cenepa South America, Peru, Amazonas | 0.41 | 230 | MVZ | none | 14 Jul 1978 | 25 | adult | *Hyloxalus sauli* | MVZ:Herp:162627 was originally identified as Hyloxalus sauli. However, range, toe disc morphology, and lack of oblique lateral line make H. sauli ID unlikely. H. bocagei and H. italoi both occur at the specimen collection site, however our specimen lacks the ventrolateral line expected in H. bocagei. Thus, coloration and range favor identification as H. italoi, and we revised the ID accordingly, though with some uncertainty. |
| *Hyloxalus lehmanni* | 1 | MVZ:Herp:192351 | 4.9166667 | -74.45 | canyon immediately N Granja Infantil, 6 km N (by road) Alban, South America, Colombia, Cundinamarca | 6.00 | 2000 | MVZ | potentially large, range of elevation in region is 1500-2500 | 14 Jul 1977 | 15.49 | juvenile | *Colostethus inguinalis* | See notes for MVZ:Herp:192350 for a comparison of characteristics of a sympatric frog. This is likely a juvenile of *Hyloxalus lehmanni.* The identification was revised with some uncertainty. |
| *Hyloxalus lehmanni* | 0 | MVZ:Herp:192350 | 4.9166667 | -74.45 | canyon immediately N Granja Infantil, 6 km N (by road) Alban, South America, Colombia, Cundinamarca | 6.00 | 2000 | MVZ | potentially large, range of elevation in region is 1500-2500 | 14 Jul 1977 | 18.85 | adult | *Colostethus inguinalis* | MVZ:Herp:192350 was originally identified as Colostethus inguinalis. However, specimen coloration makes C. inguinalis identification unlikely. Hyloxalus bocagei, Hyloxalus subpunctatus, Allobates niputaidea, and Allobates cepedai occur at the specimen locality although finger and toe pad morphology makes these indentifications unlikley. Leucostethus jota, Hyloxalus edwardsi, and Hyloxalus ruizi occur at the specimen locality but are unlikely correct idenfications due to differences in webbing morphology. Hyloxalus arliensis also occurs at the site but is this species is smaller than the specimen in concern. Allobates juanii and H. felixcoperari are sympatric, but our specimen possesses neither a dorsolateral stripe (A. juanii) nor a black venter (H. felixcoperari). Specimen coloration, body size, digit morphology, and range are most consistent with identification as Hyloxalus lehmanni, and we revised the identification accordingly, although with some uncertainty. |
| *Hyloxalus subpunctatus* | 1 | MVZ:Herp:192330 | 6 | 73.1666667 | road from Charala to Duitama, 76 km S (by road) Charala, South America, Colombia, Santander | 5.00 | 3430 | MVZ | potentially moderate, site is on slope with elevation range from 2200-3400 | 12 Jul 1977 | 15.66 | juvenile | *Hyloxalus subpunctatus* | MVZ:Herp:192330 was originally identified as *Hyloxalus subpunctatus*. *H. subpunctatus* is sympatric with the morphologically similar *H. picachos*, however digit morphology favors *H. subpunctatus* over *H. picachos*. We maintained the original identification as *H. subpunctatus*. |
| *Hyloxalus sylvaticus* | 1 | MVZ:Herp:123123 | -5.316 | -79.52617 | 6.4 mi E (by road) Canchaque, South America, Peru, Piura | 4.86 | 2590 | MVZ | potentially moderate, site is on slope with elevation range from 2000-3000 | 25 Feb 1972 | 17.06 | adult | *Hyloxalus sylvaticus* | MVZ:Herp:123123 was originally identified as *Hyloxalus sylvaticus*. It could be confused with *H. elachyhistus*, which is sympatric, but compared to the paratype (MVZ:Herp:112482) 123123 has a clearly distinct snout morphology and a different genome size, suggesting they are different species. thus, we maintained the original identification as *H. sylvaticus* with uncertainty. |
| *Colostethus ramirezi* | 1 | MVZ:Herp:192343 | 6.25 | -75.80924 | 5-6 km S (by road) San Felix on road to Microwave Station, ca. 25 km W (by road) Medellin, South America, Colombia, Antioquia | 0.50 | 2800 | MVZ | moderate, range of elevation in region is 1200-1500 | 22 Jul 1977 | 18.8 | adult | *Colostethus latinasus* | MVZ:Herp:192343 was originally identified as Colostethus latinasus, however, this species does not occur at the specimen collection site in Colombia. Several species of Colosthesus, Hyloxalus, and Allobates occur at the specimen locality. Of these species, some lack any published morphological data, so we reviewed only those with available data. Three of the four candidate Colostethus species (C. fraterdanieli, C. pratti, and C. inguinalis) as well as Hyloxalus lehmanni can be exluded on the basis of coloration. The remaining candidate Hyloxalus species and most candidate Allobates species can be excluded on the basis of digit morphology. Allobates talamancae can be excluded based on size (specimen of interest of below expected size range of A. talamancae). Range, digit morphology, coloration, and body size match most closely with that of Colostethus ramirezi, and we revised the identificaion accordingly, although with some uncertainty. |
| *Platyplectrum ornatum* | 1, 2 | MVZ:Herp:77571 | -11.96666 | 141.9 | Mapoon Mission Station, Cape York Peninsula, Australia, Australia, Queensland | 0.94 | NA | NA | none | 16 Mar 1960 | 21.26 | NA | *Limnodynastes ornatus* | MVZ:Herp:77571 was originally identified as *Lymnodynastes ornatus.* The specimen morphologically and collection site matches that of *Lymnodynastes ornatus*. We mainained the original identification as *Lymnodynastes ornatus*. The taxanomy of this species has since been updated to *Platyplectrum ornatum* |
| *Platyplectrum ornatum* | 1, 2 | MVZ:Herp:77575 | -11.96666 | 141.9 | Mapoon Mission Station, Cape York Peninsula, Australia, Australia, Queensland | 0.94 | NA | NA | none | 16 Mar 1960 | 18.96 | NA | *Limnodynastes ornatus* | MVZ:Herp:77575 was originally identified as *Lymnodynastes ornatus.* The specimen morphologically and collection site matches that of *Lymnodynastes ornatus*. We mainained the original identification as *Lymnodynastes ornatus*. The taxanomy of this species has since been updated to *Platyplectrum ornatum* |
| *Limnodynastes peronii* | 2 | MVZ:Herp:77567 | -17.53333 | 146.03333 | Innisfail [Town], Johnstone Shire, Australia, Australia, Queensland | 1.98 | NA | NA | NA | 4 Sept 1959 | 35.92 | NA | *Limnodynastes peronii* | MVZ:Herp:77567 was originally identified as *Lymnodynastes peronii.* The specimen morphologically and collection site matches that of *Lymnodynastes peronii*. We mainained the original identification as *Lymnodynastes peronii*. |
| *Limnodynastes peronii* | 2 | MVZ:Herp:77568 | -17.53333 | 146.03333 | Innisfail [Town], Johnstone Shire, Australia, Australia, Queensland | 1.98 | NA | NA | NA | 13 Dec 1959 | 31.07 | NA | *Limnodynastes peronii* | MVZ:Herp:77568 was originally identified as *Lymnodynastes peronii.* The specimen morphologically and collection site matches that of *Lymnodynastes peronii*. We mainained the original identification as *Lymnodynastes peronii*. |
| *Mannophryne trinitatis* | 1, 2 | MVZ:Herp:199838 | 10.46786 | -61.19965 | Tamana Cave, Charuma Ward, Trinidad, West Indies, Trinidad and Tobago, Saint Andrew Parish, Lesser Antilles, Trinidad | 5.15 | 266 | RM_Approx | small, elevation range in area is 0-200 | 12 Jan 1985 | 23.28 | adult | *Mannophryne trinitatus* | MVZ:Herp:199838 was originally identified as *Mannophryne trinitatis*. Morphological characters and specimen collection site match well with that of *Mannophryne trinitatis*, and no other dendrobatids occur at this location. We maintained the original identification as *Mannophryne trinitatis.* |
| *Mannophryne trinitatis* | 2 | MVZ:Herp:83764 | 10.71502 | -61.3103 | Blanchisseuse Rd., 6 mi Post, Trinidad, West Indies, Trinidad and Tobago, Saint George Parish, Lesser Antilles, Trinidad | 12.40 | 250 | RM_Approx | potentially large, elevation ranges from 0-800m in region. 6Mi from either end of the road gets to ~250m. Used that. | 20 Jul 1960 | 18.82 | adult | *Mannophryne trinitatus* | MVZ:Herp:83764 was originally identified as *Mannophryne trinitatis*. Morphological characters and specimen collection site match well with that of *Mannophryne trinitatis*, and no other dendrobatids occur at this location. We maintained the original identification as *Mannophryne trinitatis.* |
| *Oophaga pumilio* | 2 | MVZ:Herp:200894 | NA | NA | Puerto Leon [=Puerto Limon?], Central America, Costa Rica, Limon | NA | 4 | RM_Approx | likely the same locality as 110423 given the collection date, but can't confirm | 1 June 1973 | 24.8 | adult | *Oophaga pumilio* | MVZ:Herp:200894 was originally identified as *Oophaga pumilio*. *O. pumilio* is sympatric with the morphologically similar *Silverstoneia flotator, Silverstoenia nubicola,* and *Allobates talamancae*. This specimen lacks an oblique lateral strip, which excludes the aforementioned three species. *O. pumilio* is also sympatric with *O. granulifera*, however our specimen lacks the dorsal granules expected in *O. granulifera*. Range and morphological characters best support *O. pumilio*. We maintained the original identification as *Oophaga pumilio.* |
| *Oophaga pumilio* | 0 | MVZ:Herp:110423 | 9.98333 | -83.05 | 3.5 km W Puerto Limon, W edge of Pueblo Nuevo Central America, Costa Rica, Limon | 3.80 | 4 | RM_Approx | none; took elevation from Google Maps | 7 Jul 1973 | 19.89 | adult | *Oophaga pumilio* | MVZ:Herp:110423 was originally identified as *Oophaga pumilio*. *O. pumilio* is sympatric with the morphologically similar *Silverstoneia flotator, Silverstoenia nubicola,* and *Allobates talamancae*. This specimen lacks an oblique lateral strip, which excludes the aforementioned three species. *O. pumilio* is also sympatric with *O. granulifera*, however our specimen lacks the dorsal granules expected in *O. granulifera*. Range and morphological characters best support *O. pumilio*. We maintained the original identification as *Oophaga pumilio.* |
| *Phyllobates aurotaenia* | 2 | RMPC058 | NA | NA | NA | NA | NA | NA | NA | NA | 16.54 | juvenile | *Phyllobates aurotaenia* | Parents were genotyped as *P. aurotaenia.* |
| *Phyllobates aurotaenia* | 2 | RMPC059 | NA | NA | NA | NA | NA | NA | NA | NA | 30.18 | ? | *Phyllobates aurotaenia* | Parents were genotyped as *P. aurotaenia.* |
| *Phyllobates bicolor* | 2 | RMPC055 | NA | NA | NA | NA | NA | NA | NA | NA | 33.79 | juvenile | *Phyllobates bicolor* | Parents were genotyped as *P. bicolor.* |
| *Phyllobates lugubris* | 1 | MVZ:Herp:82858 | 10.3637 | -83.6349 | La Colonia, 23 km NE Guapiles, Central America, Costa Rica, Limon | 12.04 | 42 | RM_Approx | likely small, although there is a short mountain nearby, elevation vaires by ~10m on a 23Km radius around Guapiles. | 21 Aug 1966 | 18.87 | adult | *Phyllobates lugubris* | MVZ:Herp:82858 was originally identified as *Phyllobates lugubris*. It could be confused with *P. vittatus* but we retain the original identification based on the presence of a ventrolateral stripe. |
| *Phyllobates terribilis* | 2 | RMPC056 | NA | NA |  | NA | NA |  |  |  | NA | juvenile | *Phyllobates terribilis* | Parents were genotyped as *P. terribilis.* |
| *Phyllobates terribilis* | 2 | RMPC057 | NA | NA |  | NA | NA |  |  |  | 34.31 | juvenile | *Phyllobates terribilis* | Parents were genotyped as *P. terribilis.* |
| *Phyllobates vittatus* | 1 | MVZ:Herp:187229 | 8.5145 | -83.5634 | Sirena, Corcovado National Park, Central America, Costa Rica, Puntarenas | 8.30 | 126 | RM_Approx | potentially large, area goes from 0-800m. Took elev from Google Earth from coords. | 23 Feb 1984 | 23.82 | adult | *Phyllobates vittatus* | MVZ:Herp:187229 was originally identified as *Phyllobates vittatus*. Only one *Phyllobates* species occurs at the collection locality, which is *P. vittatus*. Given the prominent dorsolateral stripe and the lack of a cream dorsal band under the eye and along the flank, this species is unlikely to be *Silverstoneia,* the only other dendrobatids near the collection site*.* We maintain the original identification as *Phyllobates vittatus.* |
| *Pseudacris nigrita* | 1 | MVZ:Herp:150303 | 34.84078 | -79.38199 | 6.5 mi NE (by air) center of Laurinburg, off U.S. Rte. 401, North America, United States, North Carolina, Scotland County | 5.52 | NA | NA | none | 4 Mar 1977 | 28.37 | NA | *Pseudacris nigrita* | MVZ:Herp:150303 was originally identified as *Pseudacris nigrita*. Based on size, snout shape, white upper lip, and basal toe webbing this species does seems to be *P. nigrita*. The specimen is solid colored, and *P. nigrita* usually has a ventrolateral line but coloration seems to be very polymorphic in this species. We retained the original species identification as *P. nigrita.* |
| *Rana arvalis* | 2 | MVZ:Herp:222340 | 54.503671 | 110.694279 | Buryat Republic, Barguzin River Valley, near Kurumkan, Village Vgnasei, Asia, Russia | 0.96 | NA | NA | NA | 24 Aug 1994 | 30.15 | NA | *Rana arvalis* | Collection site within known species range, but based on locality it is possibly Rana amurensis. We retain the original identification with uncertainty. |
| *Rana cascadae* | 1, 2 | MVZ:Herp:57350 | 40.06 | -121.469 | Coon Hollow, North America, United States, California, Butte County | NA | 1524 | MVZ | NA | 3 Aug 1952 | 46.45 | NA | *Rana cascadae* | Collection site within known species range |
| *Rana cascadae* | 2 | MVZ:Herp:57348 | 40.06 | -121.469 | Coon Hollow, North America, United States, California, Butte County | NA | 1524 | MVZ | NA | 3 Aug 1952 | 53.78 | NA | *Rana cascadae* | Collection site within known species range |
| *Ranitomeya variabilis* | 1 | MVZ:Herp:174663 | -4.022 | -77.751 | vicinity of La Poza, Rio Santiago, South America, Peru, Amazonas | 1.73 | 180 | MVZ | none | 8 Aug 1979 | 16.52 | adult | *Adelphobates quinquevittatus* | MVZ:Herp:174663 as originally identified as *Adelphobates quinquevittatus*. However, it is likely a member of genus *Ranitomeya* based on size, color pattern, location, reduced digit II, and greatly expanded toe discs. It is not likely to be *R. ventrimaculata*, which has digit II = III in length according to AmphibiaWeb Ecuador. It's probably not *R. imitator*, which according to Grant char 58 should have a ventrolateral stripe; this specimen does not. Thus, it might be *R. amazonica, R. imitator, R. ventrimaculata,* or *R. variabilis*. Based on location, color pattern, and correspondence with J. Brown, the identification was revised to be R*. variabilis,* with some uncertainty*.* |
| *Ranitomeya variabilis* | 0 | MVZ:Herp:174664 | -4.022 | -77.751 | vicinity of La Poza, Rio Santiago, South America, Peru, Amazonas | 1.73 | 180 | MVZ | none | 8 Aug 1979 | 17.7 | adult | *Adelphobates quinquevittatus* | MVZ:Herp:174664 as originally identified as *Adelphobates quinquevittatus*. However, it is likely a member of genus *Ranitomeya* based on size, color pattern, location, reduced digit II, and greatly expanded toe discs. It is not likely to be *R. ventrimaculata*, which has digit II = III in length according to AmphibiaWeb Ecuador. It's probably not *R. imitator*, which according to Grant char 58 should have a ventrolateral stripe; this specimen does not. Thus, it might be *R. amazonica, R. imitator, R. ventrimaculata,* or *R. variabilis*. Based on location, color pattern, and correspondence with J. Brown, the identification was revised to be R*. variabilis,* with some uncertainty*.* |
| *Ranitomeya ventrimaculata* | 2 | MVZ:Herp:172010 | -4.0333333 | -70.1333333 | Quebrada Tucuchira, ca. 20 mi NW Leticia, South America, Colombia, Amazonas | 5.00 | NA | NA | NA | 20 Aug 1975 | 14.22 | adult | *Dendrobates sp* | MVZ:Herp:172010 was originally identified as *Dendrobates sp.* Based on its small size and coloration, it is likely a member of genus *Ranitomeya.* Only two R*anitomeya* species are found in this location, and the patterning matches *R. ventimaculata*, so we revised the identification. |
| *Rheobates palmatus* | 2 | MVZ:Herp:63210 | 4.1666667 | -73.6833333 | Buenavista, South America, Colombia, Meta | NA | 1219 | MVZ | moderate, ranges from 500 to 1200m in the region | 21 Nov 1950 | 32.66 | adult | *Rheobates palmatus* | MVZ:Herp:63210 was originally identified as *Rheobates palmatus.* Given its large size and foot webbing, it is unlikely to be any other species. The original identification was maintained. |
| *Hyloxalus sp* | 2 | MVZ:Herp:63222 | 4.1666667 | -73.6833333 | Buenavista, South America, Colombia, Meta | NA | 1219 | MVZ | moderate, ranges from 500 to 1200m in the region | 21 Nov 1950 | 28.85 | adult | *Rheobates palmatus* | EXCLUDE. It is plausibly a species of *Hyloxalus*, but given that it is male and has a large SVL, it seems to be too large to be any known *Hyloxalus* in the region. Potentially an undescribed species. It had a very different genome size estimate than for MVZ:Herp:63210, which was collected contemporaneously and in the same location. |
| *Silverstoneia erasmios* | 1 | MVZ:Herp:192358 | 3.7 | -77 | 6-8 km W Danubio on old road from Cali to Buenaventura, near La Chorrera, South America, Colombia, Valle del Cauca | 2.00 | 450 | MVZ | none | 21 Jul 1977 | 15.8 | adult | *Allobates femoralis* | MVZ:Herp:192538 was originally identified as Allobates femoralis, but based on the location and color patterning, it was revised to Silverstoneia erasmios, with some uncertainty. |
| *Silverstoneia flotator* | 1 | MVZ:Herp:113730 | 8.69745 | -83.48731 | vicinity of Tropical Science Center Field Station, Rincon de Osa, Central America, Costa Rica, Puntarenas | 1.20 | 149 | RM_Approx | none | 3 Jul 1974 | 15.65 | adult | *Silverstoneia nubicola* | MVZ:Herp:113730 was originally identified as Silverstoneia nubicola., but based on color, location, finger IV pre-axial swelling and finger IV vs V lengths, the identification was revised to S. flotator, with some uncertainty. |
| *Silverstoneia flotator* | 0 | MVZ:Herp:113729 | 8.69745 | -83.48731 | vicinity of Tropical Science Center Field Station, Rincon de Osa, Central America, Costa Rica, Puntarenas | 1.20 | 149 | RM_Approx | none | 3 Jul 1974 | 13.54 | adult | *Silverstoneia nubicola* | MVZ:Herp:113729 was originally identified as Silverstoneia nubicola., but based on color, location, finger IV pre-axial swelling and finger IV vs V lengths, the identification was revised to S. flotator, with some uncertainty. |
| *Silverstoneia nubicola* | 1 | MVZ:Herp:210478 | 9.26667 | -78.96667 | Nusagundi, Kuna Yala, Central America, Panama, Guna Yala | 11.10 | 350 | MVZ | likely small, elevation ranges 200-400 in the area | 19 Oct 1988 | 15.03 | adult | *Silverstoneia flotator* | MVZ:Herp:210478 was originally identified as Silverstoneia flotator. Based on locality, it could be *C. pratti, S. nubicola, S. flotator,* or *A. talamancae*. Based on the absence of a ventrolateral stripe, it's probably not *C. pratti*. It is small for an *Allobates*. It has a prominent dark stripe on the dorsal side of legs, which is normally absent in *S. flotator*. In addition, *S. nubicola* tends to be closer to 17mm as compared to *S. flotator* (see grant 2013). However, this individual is likely to be female (no finger IV swelling). Basal webbing between toe III and IV. character 5 (grant 2017) is state 1 (finger V reaches distal tubercle of finger IV) suggesting *S. nubicola*. The identification was revised to *S. nubicola,* with some uncertainty. |
| *Xenopus laevis* | 1, 2 | MVZ:Herp:267342 | -22.63177778 | 30.39936111 | Nwanedi Dam Camp, Africa, South Africa, Limpopo Province | 0.03 | 537 | MVZ | none | 7 Jun 2011 | NA | NA | *Xenopus laevis* | Collection site within known species range |
| *Xenopus laevis* | 0 | MVZ:Herp:234828 | NA | NA | Santchou Ville, S of Dchang, Africa, Cameroon, Sud-Ouest Province | NA | NA | NA | NA | 2 Jun 1999 | 62.55 | NA | *Xenopus laevis* | Collection site within known species range |

**Table S2**. Published c-values used in analyses and/or in our standard curve, with notes on revised species identifications. Some specimens that were not used in this study but whose identification was updated are also included.

| **Updated species identification** | **haploid c-value** | **Species identification listed in reference** | **Notes on revision of identification** | **Specimens** | **Reference** |
| --- | --- | --- | --- | --- | --- |
| *Aromobates tokuko* | 0.795 | *Aromobates tokuko* | We did not inspect the specimen, and its metadata were not available online so we were unable to verify its identity. | MNCN 59532 | Liedtke et al. 2018 |
| *Anaxyrus cognatus* | 5.6 | *Bufo cognatus* | Although there are no museum specimens to review, this species is known to occur at the collection site. | none listed; collected in near the Southwestern Research Station at Portal, Arizona | (Bachmann 1972) |
| *Lithodytes lineatus*  (excluded) | 7.11 | *Dendrobates femoralis* | in ROM database, this specimen is now *Lithodytes lineatus* | ROM 22786-22787 | (MacCulloch et al. 1996) |
| *Allobates femoralis* | 8.49 | *Epipedobates trivittatus* | Based on size and pattern of dendrobatids at collection site (pers. comm. Max Ringler and Andrius Pašukonis) | TCWC 65465 | (Camper et al. 1993) |
| *Anomaloglossus* cf. *surinamensis* | 6.9 | *Colostethus marchesianus* | We could not verify the identification of this specimen because the specimen had no remaining pigmentation. In any case, it is not *A. marchesianus,* which does not occur in French Guiana. It is also not *Allobates* *granti*, which occurs in the same area, because this specimen has a median lingual process. This means that this specimen must be a member of the genus *Anomaloglossus,* which is the only other dendrobatid group in the region. SVL was measured using photographs of the specimen. | TCWC 65478 | (Camper et al. 1993) |
| Unknown  (excluded) | 6.55 | *Epipedobates* cf. *pictus* sp. 1 | Unlikely to be *A. hahneli* or *A. femoralis* either given the size and coloration, unless they are juveniles; species unknown. No median lingual process, so not *Anomaloglossus* | ROM 22797-99 | (MacCulloch et al. 1996) |
| *Allobates* cf. *femoralis* | 8.9 | *Epipedobates* cf. *pictus* sp. 2 | Based on range, size, and pattern, either *A. hahneli* or *A. femoralis*. Likely *A. femoralis* because Finger V length is character state 1, reaching distal half of distal subarticular  tubercle of Finger IV  (*A. hahneli* is 0, surpassing tubercle), plus body shape looks like *A. femoralis* | ROM 22795-6, ROM 22805-7 | (MacCulloch et al. 1996) |
| No change | 10.30 | *Bombina bombina* | Although there are no museum specimens to review, this species is known to occur at the collection site. | Collected from unspecified USSR locality | (Mazin 1980) |
| No change | 10.57 | *Bombina bombina* | It is not possible to review the captive-bred specimens without more information. | captive bred from Zoologische Handlung | (Ullerich 1970) |
| No change | 10.92 | *Bombina bombina* | Although there are no museum specimens to review, this species or a close relative (*B. variegata*) is known to occur at the collection site. | Collected from Serbia | (Borkin et al. 2005) |
| No change | 11.85 | *Bombina bombina* | Not possible to review because specimen information was not included in the publication. | no information provided | (Vinogradov 1998) |
| No change | 12.26 | *Bombina bombina* | N/A | N/A | (Sexsmith 1971) |
| No change | 12.40 | *Bombina bombina* | Not possible to review because specimen information was not included in the publication. | no information provided | (Olmo 1973) |
| No change | 6.9 | *Bombina variegata* | It is not possible to review the captive-bred specimens without more information. | Captive-bred, origin unknown | (Horner and Macgregor 1983) |
| No change | 8.75 | *Bombina variegata* | Although there are no museum specimens to review, this species is known to occur at the collection site. | Collected from unspecified USSR locality | (Mazin 1980) |
| No change | 9.15 | *Bombina variegata* | It is not possible to review the captive-bred specimens without more information. | Captive-bred, from Germany or Israel | (Chipman et al. 2001) |
| No change | 9.17 | *Bombina variegata* | Not possible to review because specimen information was not included in the publication. | no information provided | (de Semet 1981) |
| No change | 9.6 | *Bombina variegata* | Although there are no museum specimens to review, this species is known to occur at the collection site. It is not possible to review the captive-bred specimens without more information. | Some captive-bred from Zoologische Handlung; others collected near Öhringen, Germany | (Ullerich 1970) |
| No change | 9.8 | *Bombina variegata* | N/A | N/A | (Vialli 1957) |
| No change | 8.95 | *Dendrobates tinctorius* | We reviewed photographs of the specimens and determined that the identification is correct based on size, pattern, and collection locality. SVL was measured using photographs of the specimen. | TCWC 65473-65476 | (Camper et al. 1993) |
| *Ctenophryne geayi* (excluded) | 2.04 | *Elachistocleis ovalis* | This specimen was reidentified by MacCulloch in 2001 as *Ctenophryne geayi* (see vertnet entry). It is not clear why or how the specimen was reidentified, but it might be because *E. ovalis* is considered a *nomen dubium* (Barrio-Amorós et al. 2019) | ROM 22769 | (MacCulloch et al. 1996) |
| Unknown  (excluded) | 5.39 | *Elachistocleis ovale* | “*Elachistocleis ovalis”* is a *nomen inquirenda,* so animals are being reidentified. This animal is from Cayenne, French Guiana. | TCWC 65482 | (Camper et al. 1993) |
| No change | 9.20 | *Hyla versicolor* | Not possible to review because specimen information was not included in the publication. | no information provided | (Vinogradov 1998) |
| No change | 10.30 | *Hyla versicolor* | Although there are no museum specimens to review, this species is known to occur at the collection site. | specimen is from an unspecified site in the USA | (Bachmann and Nishioka 1978) |
| *Platypectrum ornatum* | 0.95 | *Limnodynastes ornatus* | Not possible to review because specimen information was not included in the publication. | no information provided | (Heyer and Liem 1976; Olmo and Morescalchi 1978) |
| No change | 1.3 | *Limnodynastes peronii* | Not possible to review because specimen information was not included in the publication. | no information provided | (Olmo and Morescalchi 1978) |
| No change | 2.25 | *Limnodynastes peronii* | Not possible to review because specimen information was not included in the publication. | no information provided | (Olmo 1973) |
| No change | 3.08 | *Limnodynastes peronii* | Not possible to review because specimen information was not included in the publication. | no information provided | (Vinogradov 1998) |
| No change | 2.98 | *Mannophryne trinitatis* | Not possible to review because specimen information was not included in the publication. However, we are confident in this identification considering *M. trinitatis* is endemic to the island of Trinidad, and the only species in the subfamily Aromobatinae on the island. | none listed | (Camper et al. 1993) |
| No change | 4.689 | *Oophaga pumilio* | Based on the collection locality we are confident in this identification. Unable to review specimens as they are not currently held at the MVZ. | MVZ:Herp:269267-269268 | (Liedtke et al. 2018) |
| No change | 4.05 | *Pseudacris nigrita* | Although it is not clearly described in the article, it seems like these data were taken from a specimen collected in Florida, which does overlap with the known range of this species. | specimens collected in Florida | (Goin et al. 1968) |
| No change | 4.65 | *Rana arvalis* | It is not possible to review the captive-bred specimens without more information. | Captive-bred, origin unknown (but likely Germany) | (Ullerich 1967) |
| No change | 5.15 | *Rana arvalis* | Although there are no museum specimens to review, this species is known to occur at the collection site. | Collected from unspecified USSR locality | (Mazin 1980) |
| No change | 5.23 | *Rana arvalis* | Not possible to review because specimen information was not included in the publication. | no information provided | (Ullerich 1970) |
| No change | 6.18 | *Rana arvalis* | Not possible to review because specimen information was not included in the publication. | no information provided | (Vinogradov 1998) |
| No change | 6.80 | *Rana arvalis* | Although there are no museum specimens to review, this species is known to occur at the collection site. | collected in East Germany | (Oeldorf et al. 1978) |
| No change | 7.14 | *Rana arvalis* | Not possible to review because specimen information was not included in the publication. | no information provided | (Olmo 1973) |
| No change | 7.17 | *Rana arvalis* | N/A | N/A | (Sexsmith 1971) |
| No change | 7.49 | *Rana cascadae* | Not possible to review because specimen information was not included in the publication. | no information provided | (Olmo 1973) |
| No change | 7.91 | *Rana cascadae* | N/A | N/A | (Sexsmith 1971) |
| No change | 8.63 | *Rana cascadae* | Not possible to review because specimen information was not included in the publication. | no information provided | (Vinogradov 1998) |
| No change | 3.00 | *Xenopus laevis* | It is not possible to review the captive-bred specimens without more information. | Captive-bred, origin unknown | (Horner and Macgregor 1983) |
| No change | 3.09 | *Xenopus laevis* | Not possible to review because specimen information was not included in the publication. | no information provided | (Olmo 1973) |
| No change | 3.11 | *Xenopus laevis* | Although there are no museum specimens to review, this species is known to occur at the collection site. | collected in Fish Hock, Cape, South Africa | (Thiébaud and Fischberg 1977) |
| No change | 3.15 | *Xenopus laevis* | Not possible to review because specimen information was not included in the publication. | no information provided | (Dawid 1965) |
| No change | 3.15 | *Xenopus laevis* | Not possible to review because specimen information was not included in the publication. | either captive-bred or collected at an unspecified site in Africa | (Giorgi and Fischberg 1982) |
| No change | 3.59 | *Xenopus laevis* | Not possible to review because specimen information was not included in the publication. | no information provided | (Vinogradov 1998) |
| No change | 3.69 | *Xenopus laevis* | N/A | N/A | (Sexsmith 1971) |
| No change | 3.85 | *Xenopus laevis* | Not possible to review because specimen information was not included in the publication. | no information provided | (de Semet 1981) |

**Table S3**. GenBank numbers for sequences used to time-calibrate the phylogeny used in analyses.

| **Species** | **Grant2017_Taxon_label** | **H1_ND2** | **RHO** | **TYR** | **RAG1** | **POMC** | **SLC8A1** | **Genome Size Locality** | **Grant et al. 2017 Locality** |
| --- | --- | --- | --- | --- | --- | --- | --- | --- | --- |
| *Allobates femoralis* | Allobates_femoralis_femo04_Yurimaguas | DQ523072 | - | - | - | - | - | 2 mi inland from Bella Vista on Caijaru River, South America, Peru, Loreto | Peru: Loreto: Yurimaguas: Shucshuyacu |
| *Allobates talamancae* | Allobates_talamancae_SIUC7667 | DQ502166 | DQ503250 | - | DQ503363 | - | - | vicinity of Tropical Science Center Field Station, Rincon de Osa, Central America, Costa Rica, Puntarenas | Panama: Coclé: El Copé, Parque Nacional General de División "Omar Torrijos Herrera" |
| *RoberAmeerega hahneli* | Ameerega_hahneli_Leticia_ICN50410 | DQ502270 | DQ503276 | DQ503174 | - | - | - | 46 km S and 22 km W San Martin, South America, Colombia, Meta | Colombia: Amazonas: Leticia, Lago Yahuarcaca |
| *Ameerega parvula* | Ameerega_parvula_QCAZ16584 | HQ290999 | - | HQ290936 | - | HQ290876 | HQ290756 | vicinity of Huampami (Aguaruna village), Rio Cenepa South America, Peru, Amazonas | Ecuador: Morona Santiago: near M_ndez, 550 m |
| *Ameerega trivittata* | Ameerega_trivittata_MPEG12504 | DQ502079 | DQ503211 | DQ503152 | DQ503324 | - | - | Mazaroni Rd., 0.3 km from junction with Brownsberg Rd., Brownsberg Nature Park, South America, Suriname, Brokopondo District | Brazil: Acre: Porto Walter, 8°5'31.2"S, 72°46'37.1"W |
| *Andinobates minutus* | Andinobates_minutus_KRL790 | DQ502168 | DQ503251 | - | DQ503365 | - | - | Nusagundi, Kuna Yala, Central America, Panama | Panama: Coclé: El Copé, Parque Nacional General de División "Omar Torrijos Herrera" |
| *Anomaloglossus stepheni* | Anomaloglossus_stepheni_MJH3928 | DQ502107 | DQ503223 | - | DQ503334 | - | - | Brownsberg Nature Park Headquarters, South America, Suriname, Brokopondo District | Brazil: Amazonas: Reserva Florestal Adolfo Ducke |
| *Colostethus panamansis* | Colostethus_panamansis_CH5546 | DQ502172 | - | - | DQ503368 | - | - | 1 km SW (by air) Hotel Greco, El Valle de Anton, Central America, Panama, Cocle | Panama: Darién: Caná, 1246 m |
| *Dendrobates auratus* | Dendrobates_auratus_USNM313818 | AY843581 | AY844554 | AY844032 | DQ503304 | - | - | Isla Taboga Pacific Ocean, Gulf of Panama, Panama, Panama, Isla Taboga | Panama: Bocas del Toro: Laguna de Tierra Oscura, 3.7 km S of Tiger Key |
| *Dendrobates leucomelas* | Dendrobates_leucomelas_TNHCFS5639 | HQ290987 | - | HQ290924 | - | HQ290864 | HQ290744 | Ucaima, South America, Venezuela, Bolivar | Venezuela: Amazonas: Puerto Ayacucho, Tobogán, 81 m |
| *Dendrobates truncatus* | Dendrobates_truncatus_TNHCFS4979 | HQ290992 | - | HQ290929 | - | HQ290869 | HQ290749 | 8 km N Chaparral, South America, Colombia, Tolima | Colombia: Tolima: Mariquita, vereda Malabares, 587 m |
| *Epipedobates boulengeri* | Epipedobates_boulengeri_QCAZ16574 | EU342569 | - | HQ290934 | - | HQ290874 | HQ290754 | 6-8 km W Danubio on old road from Cali to Buenaventura, near La Chorrera, South America, Colombia, Valle del Cauca | Ecuador: Esmeraldas: A 3 Km de Durango, road to San Lorenzo, 253 m |
| *Hyloxalus elachyhistus* | Hyloxalus_elachyhistus_QCAZ16517 | HQ290956 | - | HQ290896 | - | HQ290836 | HQ290716 | 2 km N Loja, South America, Ecuador, Loja | Ecuador: El Oro: Torata-Balsas road |
| *Hyloxalus infraguttatus* | Hyloxalus_infraguttatus_QCAZ16516 | AY364548 | - | - | - | - | - | 95 km E Guayaquil on road to Quito, South America, Ecuador, Guayas | Ecuador: Manabí: road Portoviejo -Jipijapa at 6 Km from Puerto Cayo, -80.708 -1.324 |
| *Hyloxalus italoi* | Hyloxalus_italoi_QCAZ16511 | AY364558 | - | - | - | - | - | vicinity of Huampami (Aguaruna village), Rio Cenepa, South America, Peru, Amazonas | Ecuador: Morona Santiago: Santiago |
| *Hyloxalus lehmanni* | Hyloxalus_lehmanni_MAR2675 | MF624228 | - | - | - | - | - | canyon immediately N Granja Infantil, 6 km N (by road) Alban, South America, Colombia, Cundinamarca | Colombia: Tolima: Río Blanco, corregimiento de Herrera, Las Mercedes Parte Alta (ca de 12.5 Km al SW de Herrera), 2450 m. 3°17’3.3’’N, 75°55’6.3’’W |
| *Hyloxalus subpunctatus* | Hyloxalus_subpunctatus_TNHCFS4957 | HQ290973 | - | HQ290911 | - | HQ290851 | HQ290731 | road from Charala to Duitama, 76 km S (by road) Charala, South America, Colombia, Santander | Colombia: Boyacá: Chiquinquira, 2575 m |
| *Hyloxalus sylvaticus* | Hyloxalus_sylvaticus_KU219756 | DQ501988 | DQ503178 | DQ503141 | DQ503286 | - | - | 6.4 mi E (by road) Canchaque, South America, Peru, Piura | Peru: Piura: Ayacaba, 12.7 km E El Tambo, 2820 m |
| *Colostethus ramirezi* | Colostethus_ramirezi_MAR3312 | MF624216 | MF614406 | MF624158 | - | - | - | 5-6 km S (by road) San Felix on road to Microwave Station, ca. 25 km W (by road) Medellin, South America, Colombia, Antioquia | Colombia: Antioquia, Urrao: zona de amortiguación Parque Nacional Natura Las Orquídeas, "Páramo El Almorzadero", 2160 m |
| *Mannophryne trinitatis* | Mannophryne_trinitatis_MVZ199828 | DQ502131 | DQ503236 | - | DQ503345 | - | - | Tamana Cave, Charuma Ward, Trinidad, West Indies, Trinidad and Tobago, Saint Andrew Parish, Lesser Antilles, Trinidad AND Blanchisseuse Rd., 6 mi Post, Trinidad, West Indies, Trinidad and Tobago, Saint George Parish, Lesser Antilles, Trinidad | Trinidad and Tobago: Nariva Parish, Charuma Ward, Tamana Cave |
| *Oophaga pumilio* | Oophaga_pumilio_TNHCFS4814 | HQ290988 | - | HQ290925 | - | HQ290865 | HQ290745 | Puerto Leon [=Puerto Limon?], Central America, Costa Rica, Limon | Panama: Bocas del Toro: Isla Colón, Bocas del Drago (Dragomar), 11 m |
| *Phyllobates aurotaenia* | Phyllobates_aurotaenia_TNHCFS4990 | HQ291005 | - | HQ290942 | - | HQ290882 | HQ290762 | Captive bred | Colombia: Chocó: Quibdó road to Pacuritas, 50 m |
| *Phyllobates bicolor* | Phyllobates_bicolor_1233 | DQ502181 | - | 0 | DQ503377 | - | - | Captive bred | No data (captive bred) |
| *Phyllobates lugubris* | Phyllobates_lugubris_USNMFS195116 | DQ283043 | DQ503193 | - | DQ503306 | - | - | La Colonia, 23 km NE Guapiles, Central America, Costa Rica, Limon | Panama: Bocas del Toro: S end of Isla Popa, 1 km E Sumwood Channel |
| *Phyllobates terribilis* | Phyllobates_terribilis_AMNHA118566 | DQ502157 | DQ503244 | - | DQ503358 | - | - | Captive bred | Colombia: Cauca: Quebrada Guanguí, 0.5 km above Rio Patia (upper Saija drainage), 100-200 m |
| *Phyllobates vittatus* | Phyllobates_vitattus_839 | DQ502152 | - | DQ503166 | DQ503355 | - | - | Sirena, Corcovado National Park, Central America, Costa Rica, Puntarenas | No data (captive bred) |
| *Ranitomeya variabilis* | Ranitomeya_variabilis_Sucumbios_OMNH34091 | DQ502069 | DQ503205 | - | DQ503319 | - | - | vicinity of La Poza, Rio Santiago, South America, Peru, Amazonas | Ecuador: Sucumbíos: Estación Científica de Universidad Católica near Reserva Faunística Cuyabeno, 220 m, 0°0'S 76°10'W |
| *Ranitomeya ventrimaculata* | Ranitomeya_ventrimaculata_JDL24489 | DQ502266 | DQ503272 | - | DQ503399 | - | - | Quebrada Tucuchira, ca. 20 mi NW Leticia, South America, Colombia, Amazonas | Colombia: Amazonas: Leticia, Km 11 (Leticia-Tarapacá) |
| *Rheobates palmatus* | Rheobates_palmatus_MUJ3829 | MF624243 | MF614424 | - | - | - | - | Buenavista, South America, Colombia, Meta | COLOMBIA: Meta: Villavicencio, Vereda El Carmen, Caño Blanco, 900_1000 m |
| *Silverstoneia erasmios* | Silverstoneia_erasmios_LSB218 | MF624244 | MF614425 | MF624169 | MF614372 | - | - | 6-8 km W Danubio on old road from Cali to Buenaventura, near La Chorrera, South America, Colombia, Valle del Cauca | COLOMBIA: Antioquia: Frontino, Murrí-La Blanquita, 500 m |
| *Silverstoneia flotator* | Silverstoneia_flotator_TNHCFS4804 | HQ290957 | - | HQ290897 | - | HQ290837 | HQ290717 | vicinity of Tropical Science Center Field Station, Rincon de Osa, Central America, Costa Rica, Puntarenas | Panama: Coclé: El Copé, Parque Nacional General de División Omar Torrijos Herrera, 782 m |
| *Silverstoneia nubicola* | Silverstoneia_nubicola_SIUC7652 | DQ502161 | DQ503245 | - | DQ503359 | - | - | Nusagundi, Kuna Yala, Central America, Panama, Guna Yala | Panama: Coclé: El Copé, Parque Nacional General de División "Omar Torrijos Herrera" |
| *Anomaloglossus cf_surinamensis* | NA | KY510066 | - | - | KY549451 | KY549494 | - | French Guiana, Cayenne: 65 km W Iracoubo | French Guiana: Fleuve Mana, 5.0994 -53.8005 |
| *Dendrobates tinctorius* | Dendrobates_tinctorius_UTAA56495 | DQ502248 | DQ503266 | - | DQ503387 | - | - | French Guiana, Cayenne: Kaw Mts, I 50 km SE Cayenne | Suriname: Sipaliwini: ca. 1.0 km N of Tafelberg airstrip |

**Table S4.** Morphological, life history, and metabolic data used in regression analyses models. Life history data were obtained from Carvajal-Castro et al. (2021), metabolic and body weight data were obtained from Santos (2012) and Santos and Cannatella (2011), alkaloid data from Santos et al. (2016), and morphological data were measured on museum specimens (see Table S1), or obtained from the literature (see Methods for details).

| **Species** | **Mean c-value** | **Habit (binary)** | **Breeding site (binary)** | **Mean number of eggs** | **Mean clutch size** | **Body Mass (g)** | **RMR (VO_2_ ml*h^-1^)** | **AMR (VO_2_ ml*h^-1^)** | **Scope (raw)** | **Embryo time (days)** | **Time to metamorphosis (days)** | **Adult age (months)** | **Longevity (years)** | **Alkaloids (imputed)** | **SVL (mm)** | **Elevation (masl)** | **Cell size (μm)** |
| --- | --- | --- | --- | --- | --- | --- | --- | --- | --- | --- | --- | --- | --- | --- | --- | --- | --- |
| *Allobates femoralis* | 8.49 | terrestrial | Phytotelmata | NA | 28 | 1.246 | 0.223 | 1.104 | 0.881 | NA | 45 | 9 | 6 | 0 | 23.93 | 600 | 738.49 |
| *Allobates talamancae* | 7.30 | terrestrial | Stream | NA | 34 | 0.88 | 0.08 | 0.714 | 0.643 | 17 | 77.5 | 12 | NA | 0 | 20.225 | 410 | 756.42 |
| *Ameerega hahneli* | 5.97 | terrestrial | Stream | NA | 22 | 0.343 | 0.1 | 0.516 | 0.416 | NA | 60 | 8.5 | 3 | 1 | 21.765 | 488 | 728.31 |
| *Ameerega parvula* | 4.51 | terrestrial | Stream | 18.4 | 10 | 1.569 | 0.293 | 1.497 | 1.204 | NA | 60 | 12 | NA | 1 | 26.295 | 213 | 779.62 |
| *Ameerega trivittata* | 7.62 | terrestrial | Stream | 32.5 | 38.4 | 5.512 | 0.964 | 6.632 | 5.668 | 18.5 | 65 | NA | 6 | 1 | 35.825 | 495 | 823.17 |
| *Anaxyrus cognatus* | 5.60 | NA | NA | NA | NA | NA | NA | NA | NA | NA | NA | NA | NA | NA | 23.99 | NA | NA |
| *Andinobates minutus* | 3.29 | terrestrial | Phytotelmata | 2 | 2 | 0.16 | 0.051 | 0.577 | 0.526 | 14 | NA | NA | NA | 1 | 13.32 | 250 | 881.15 |
| *Anomaloglossus stepheni* | 3.52 | terrestrial | NA | NA | 3.57 | NA | NA | NA | NA | NA | NA | NA | NA | 0 | 19.19 | 495 | 783.67 |
| *Anomaloglossus surinamensis* | 6.90 | terrestrial | NA | NA | 2 | NA | NA | NA | NA | NA | NA | NA | NA | 0 | 15.625 | NA | NA |
| *Bombina bombina* | 11.38 | NA | NA | NA | NA | NA | NA | NA | NA | NA | NA | NA | NA | NA | NA | NA | NA |
| *Bombina variegata* | 8.90 | NA | NA | NA | NA | NA | NA | NA | NA | NA | NA | NA | NA | NA | NA | NA | NA |
| *Colostethus panamansis* | 8.99 | riparian | Stream | NA | 25 | 0.973 | 0.096 | 1.159 | 1.063 | NA | NA | NA | NA | 1 | 25.78 | 630 | 962.08 |
| *Colostethus ramirezi* | 6.14 | terrestrial | NA | NA | NA | NA | NA | NA | NA | NA | NA | NA | NA | 0 | 18.8 | 2800 | 833.53 |
| *Dendrobates auratus* | 5.25 | terrestrial | Phytotelmata | 9 | 7 | 1.996 | 0.306 | 2.81 | 2.504 | 16 | 87.5 | 10 | 13.25 | 1 | 22.68 | 0 | 826.11 |
| *Dendrobates leucomelas* | 4.73 | terrestrial | Phytotelmata | 5 | 5 | 2.243 | 0.414 | 3.167 | 2.753 | 13.5 | 87.5 | 13 | 10.75 | 1 | 31.34 | 427 | 797.90 |
| *Dendrobates tinctorius* | 8.95 | terrestrial | Phytotelmata | 9 | 6 | 5.037 | 0.549 | 5.983 | 5.434 | 9 | 110 | NA | 11 | 1 | 36.5 | 350 | NA |
| *Dendrobates truncatus* | 5.20 | terrestrial | Phytotelmata | 5 | 3.6 | 1.573 | 0.188 | 1.926 | 1.738 | 14 | NA | NA | NA | 1 | 28.35 | 700 | 940.04 |
| *Epipedobates boulengeri* | 5.41 | terrestrial | Stream | 20 | 20 | 0.457 | 0.097 | 0.329 | 0.232 | 15.5 | NA | 12 | 4 | 1 | 12.79 | 450 | 478.12 |
| *Hyla versicolor* | 9.75 | NA | NA | NA | NA | NA | NA | NA | NA | NA | NA | NA | NA | NA | NA | NA | 930.80 |
| *Hyloxalus elachyhistus* | 8.11 | terrestrial | Stream | NA | 13.5 | 0.958 | 0.248 | 0.84 | 0.592 | NA | NA | NA | NA | 0 | 17.235 | 2100 | 653.05 |
| *Hyloxalus infraguttatus* | 2.60 | riparian | Stream | NA | 17 | 0.806 | NA | NA | NA | NA | NA | NA | NA | 0 | 19.21 | 3000 | 524.82 |
| *Hyloxalus italoi* | 7.91 | riparian | Stream | 21 | NA | NA | NA | NA | NA | NA | NA | NA | NA | 0 | 25 | 230 | 717.29 |
| *Hyloxalus lehmanni* | 5.18 | terrestrial | Stream | NA | NA | NA | NA | NA | NA | NA | NA | NA | NA | 0 | 18.85 | 2000 | 866.78 |
| *Hyloxalus subpunctatus* | 5.35 | terrestrial | Stream | 24.5 | 21.5 | 0.799 | 0.151 | 0.901 | 0.75 | NA | NA | NA | NA | 0 | 23.6 | 3015 | 971.31 |
| *Hyloxalus sylvaticus* | 3.98 | riparian | Stream | NA | NA | NA | NA | NA | NA | NA | NA | NA | NA | 0 | 17.06 | 2590 | 673.43 |
| *Limnodynastes peronii* | 2.21 | NA | NA | NA | NA | NA | NA | NA | NA | NA | NA | NA | NA | NA | NA | NA | NA |
| *Mannophryne trinitatis* | 2.98 | riparian | Stream | 10.1875 | 13 | NA | NA | NA | NA | NA | NA | NA | NA | 0 | 21.05 | 258 | 561.36 |
| *Oophaga pumilio* | 4.25 | terrestrial | Phytotelmata | 6 | 10.9 | 0.538 | 0.067 | 0.698 | 0.631 | 12 | 47.5 | NA | 5.75 | 1 | 22.345 | 4 | 456.90 |
| *Phyllobates aurotaenia* | 10.94 | terrestrial | Stream | 30 | 12 | 2.017 | 0.184 | 1.347 | 1.163 | 14.8 | 47.5 | 12 | NA | 1 | 27.19 | NA | 1323.77 |
| *Phyllobates bicolor* | 13.00 | terrestrial | Stream | NA | 17 | NA | NA | NA | NA | 17.9 | 46.2 | NA | NA | 1 | 37.19 | NA | 1786.66 |
| *Phyllobates lugubris* | 5.04 | terrestrial | Stream | 17.5 | 15 | NA | NA | NA | NA | NA | NA | NA | NA | 1 | 18.87 | 42 | 878.17 |
| *Phyllobates terribilis* | 12.80 | terrestrial | Stream | 14 | 20 | 6.013 | 0.787 | 5.092 | 4.305 | 15 | 54.3 | 15 | 12 | 1 | 39.69 | NA | NA |
| *Phyllobates vittatus* | 11.67 | terrestrial | Stream | 14 | 15 | NA | NA | NA | NA | 19 | 51.8 | 9 | NA | 1 | 23.82 | 126 | 1024.02 |
| *Platyplectrum ornatum* | 0.95 | NA | NA | NA | NA | NA | NA | NA | NA | NA | NA | NA | NA | NA | NA | NA | NA |
| *Pseudacris nigrita* | 4.05 | NA | NA | NA | NA | NA | NA | NA | NA | NA | NA | NA | NA | NA | NA | NA | NA |
| *Rana arvalis* | 6.05 | NA | NA | NA | NA | NA | NA | NA | NA | NA | NA | NA | NA | NA | NA | NA | NA |
| *Rana cascadae* | 8.01 | NA | NA | NA | NA | NA | NA | NA | NA | NA | NA | NA | NA | NA | NA | 1524 | NA |
| *Ranitomeya variabilis* | 5.08 | terrestrial | Phytotelmata | 4 | 3.7 | NA | NA | NA | NA | NA | NA | NA | NA | 1 | 17.11 | 180 | 737.43 |
| *Ranitomeya ventrimaculata* | 4.36 | terrestrial | Phytotelmata | 4 | 2.93 | 0.346 | 0.134 | 0.661 | 0.527 | NA | 6.75 | NA | 3 | 1 | 14.22 | 350 | 756.67 |
| *Rheobates palmatus* | 6.09 | riparian | Stream | NA | 23.5 | 1.78 | 0.368 | 1.563 | 1.195 | 20 | NA | 4.5 | NA | 0 | 30.755 | 1219 | 624.42 |
| *Silverstoneia erasmios* | 3.38 | terrestrial | Stream | NA | NA | NA | NA | NA | NA | NA | NA | NA | NA | 0 | 15.8 | 450 | NA |
| *Silverstoneia flotator* | 5.17 | terrestrial | Stream | NA | 6 | 0.327 | 0.054 | 0.491 | 0.437 | NA | NA | NA | NA | 0 | 14.595 | 149 | 636.73 |
| *Silverstoneia nubicola* | 3.72 | terrestrial | Stream | NA | 6 | 0.371 | 0.043 | 0.636 | 0.593 | NA | NA | 12 | NA | 0 | 15.03 | 350 | 622.40 |
| *Xenopus laevis* | 3.33 | NA | NA | NA | NA | NA | NA | NA | NA | NA | NA | NA | NA | NA | NA | 537 | NA |

**Table S5.** Genome size estimates for each species obtained using a standard curve estimated using a linear mixed-effects model with genome size as the dependent variable, integrated optical density as a fixed-effect predictor, and sampling batch as a random-effect predictor (see Methods and Fig. 1).

| **Species** | **Median IOD (batch 1)** | **Median IOD (batch 2)** | **C-value estimate (batch 1)** | **C-value estimate (batch 2)** | **Average C-value (using IOD)** | **Other C-value measurement** | **Method (source) of other C-value measurement** | **C-value used in analyses** |
| --- | --- | --- | --- | --- | --- | --- | --- | --- |
| *Allobates femoralis* | NA | 1970456 | NA | 7.137766 | 7.137766 | 8.49 | Flow cytometry (Camper et al. 1993) | 8.49 |
| *Allobates talamancae* | NA | 2017325.5 | NA | 7.3015582 | 7.3015582 | NA | NA | 7.3015582 |
| *Ameerega hahneli* (average) | 1744137.75 | 1525835 | 6.3468643 | 5.583974 | 5.9654191 | NA | NA | 5.9654191 |
| *Ameerega hahneli* (MVZ:Herp:63120) | 1326913 | 1525835 | 4.888812 | 5.583974 | 5.236393 | NA | NA | NA |
| *Ameerega hahneli* (MVZ:Herp:63759) | 2161362 | NA | 7.804916 | NA | 7.804916 | NA | NA | NA |
| *Ameerega parvula* | 1217958 | NA | 4.5080535 | NA | 4.5080535 | NA | NA | 4.5080535 |
| *Ameerega trivittata* | 2108164.25 | NA | 7.6190072 | NA | 7.6190072 | NA | NA | 7.6190072 |
| *Anaxyrus cognatus* | NA | 1885171 | NA | 6.8397253 | 6.8397253 | 5.6 | Feulgen staining (Bachmann 1972) | 5.6 |
| *Andinobates minutus* | NA | 868470 | NA | 3.2867175 | 3.2867175 | NA | NA | 3.2867175 |
| *Anomaloglossus stepheni* | NA | 934248 | NA | 3.5165882 | 3.5165882 | NA | NA | 3.5165882 |
| *Anomaloglossus surinamensis* | NA | NA | NA | NA | NA | 6.9 | Flow cytometry (Camper et al. 1993) | 6.9 |
| *Bombina bombina* | NA | 3309447.5 | NA | 11.817064 | 11.817064 | 11.3833333 | averaged across several studies, see Table S2 | 11.383333 |
| *Bombina variegata* | NA | 2242083.5 | NA | 8.0870073 | 8.0870073 | 8.895 | averaged across several studies, see Table S2 | 8.895 |
| *Colostethus panamansis* | 2499166.25 | NA | 8.9854199 | NA | 8.9854199 | NA | NA | 8.9854199 |
| *Colostethus ramirezi* | 1686109 | NA | 6.1440745 | NA | 6.1440745 | NA | NA | 6.1440745 |
| *Dendrobates auratus* | 1429391 | NA | 5.2469365 | NA | 5.2469365 | NA | NA | 5.2469365 |
| *Dendrobates leucomelas* | 1280311.5 | NA | 4.7259568 | NA | 4.7259568 | NA | NA | 4.7259568 |
| *Dendrobates tinctorius* | NA | NA | NA | NA | NA | 8.95 | Flow cytometry (Camper et al. 1993) | 8.95 |
| *Dendrobates truncatus* | 1416045 | NA | 5.200297 | NA | 5.200297 | NA | NA | 5.200297 |
| *Epipedobates boulengeri* | 1475788.5 | NA | 5.4090793 | NA | 5.4090793 | NA | NA | 5.4090793 |
| *Hyla versicolor* | 2414754 | 2427062 | 8.6904292 | 8.7334413 | 8.7119352 | 9.75 | averaged between two studies, see Table S2 | 9.75 |
| *Hyloxalus elachyhistus* | 2248928 | NA | 8.1109264 | NA | 8.1109264 | NA | NA | 8.1109264 |
| *Hyloxalus infraguttatus* | 671580 | NA | 2.5986571 | NA | 2.5986571 | NA | NA | 2.5986571 |
| *Hyloxalus italoi* | 2190320 | NA | 7.9061123 | NA | 7.9061123 | NA | NA | 7.9061123 |
| *Hyloxalus lehmanni* | 1409342 | NA | 5.1768724 | NA | 5.1768724 | NA | NA | 5.1768724 |
| *Hyloxalus subpunctatus* | 1460145.5 | NA | 5.3544126 | NA | 5.3544126 | NA | NA | 5.3544126 |
| *Hyloxalus sylvaticus* | 1068137.5 | NA | 3.9844843 | NA | 3.9844843 | NA | NA | 3.9844843 |
| *Limnodynastes ornatus* | 467302.75 | 434227 | 1.8847809 | 1.7691929 | 1.8269869 | 0.95 | Feulgen staining (Heyer and Liem 1976) | 0.95 |
| *Limnodynastes peronii* | NA | 506315.5 | NA | 2.0211165 | 2.0211165 | 2.21 | averaged across several studies, see Table S2 | 2.21 |
| *Mannophryne trinitatis* | 925321 | 762649.75 | 3.4853915 | 2.9169134 | 3.2011524 | 2.98 | Flow cytometry (Camper et al. 1993) | 2.98 |
| *Oophaga pumilio* | 999351 | 1290909 | 3.7441 | 4.7629913 | 4.2535456 | NA | NA | 4.2535456 |
| *Phyllobates aurotaenia* | NA | 3255435.75 | NA | 11.628312 | 11.628312 | 10.9442 | Flow cytometry (this study) | 10.9442 |
| *Phyllobates bicolor* | NA | 4071766 | NA | 14.481096 | 14.481096 | 12.999 | Flow cytometry (this study) | 12.999 |
| *Phyllobates lugubris* | 1369821 | NA | 5.0387606 | NA | 5.0387606 | NA | NA | 5.0387606 |
| *Phyllobates terribilis* | NA | NA | NA | NA | NA | 12.80475 | Flow cytometry (this study) | 12.80475 |
| *Phyllobates vittatus* | NA | NA | NA | NA | NA | 11.6746 | Flow cytometry (this study) | 11.6746 |
| *Pseudacris nigrita* | NA | NA | NA | NA | NA | 4.05 | Feulgen staining (Goin et al. 1968) | 4.05 |
| *Rana arvalis* | NA | 1586563 | NA | 5.7961967 | 6.0457143 | 6.04571429 | averaged across several studies, see Table S2 | 6.0457143 |
| *Rana cascadae* | 2029612 | 1764905 | 7.3444951 | 6.4194384 | 6.8819668 | 8.01 | averaged across several studies, see Table S2 | 8.01 |
| *Ranitomeya variabilis* | 1382097 | NA | 5.0816609 | NA | 5.0816609 | NA | NA | 5.0816609 |
| *Ranitomeya ventrimaculata* | NA | 1176218 | NA | 4.3621871 | 4.3621871 | NA | NA | 4.3621871 |
| *Rheobates palmatus* (MVZ:Herp:63210) | NA | 1671435 | NA | 6.0927941 | 6.0927941 | NA | NA | 6.0927941 |
| *Rheobates palmatus* (MVZ:Herp:63222) | NA | 777832.5 | NA | 2.9699717 | 2.9699717 | NA | NA | NA |
| *Silverstoneia erasmios* | 896096 | NA | 3.3832605 | NA | 3.3832605 | NA | NA | 3.3832605 |
| *Silverstoneia flotator* | 1407624 | NA | 5.1708686 | NA | 5.1708686 | NA | NA | 5.1708686 |
| *Silverstoneia nubicola* | 992488 | NA | 3.7201162 | NA | 3.7201162 | NA | NA | 3.7201162 |
| *Xenopus laevis* | 1027238.5 | 989829 | 3.8415568 | 3.710824 | 3.7761904 | 3.32875 | averaged across several studies, see Table S2 | 3.32875 |

**Table S6.** Phylogenetically aware linear regression coefficients of genome size against 15 morphological, metabolic, and life history traits, including toxicity as a correlate. Body mass was included as a covariate in models involving metabolic rates.

| **Model** | **n** | **Adjusted r2** | **Coefficient** | **SE** | **t** | **p-value** |
| --- | --- | --- | --- | --- | --- | --- |
| log(SVL) ~ Alkaloid presence | 34 | 0.003 | 0.130 | 0.124 | 1.048 | 0.302 |
| log(SVL) ~ log(cell size) | 30 | 0.208 | 0.532 | 0.181 | 2.939 | 0.007 |
| log(clutch size) ~ log(SVL) | 29 | 0.096 | 0.808 | 0.406 | 1.991 | 0.057 |
| log(oocytes per female) ~ log(SVL) | 18 | -0.061 | 0.054 | 0.466 | 0.116 | 0.909 |
| log(Scope) ~ Alkaloid presence + | 21 | 0.819 | 0.245 | 0.144 | 0.133 | 0.200 |
| log(Mass) |  |  | 0.792 | 0.085 | 9.276 | 2.80E-08 |
| log(AMR) ~ Alkaloid presence + | 21 | 0.859 | 0.234 | 0.156 | 1.496 | 0.152 |
| log(Mass) |  |  | 0.787 | 0.074 | 10.692 | 3.16E-09 |
| log(RMR) ~ Alkaloid presence + | 21 | 0.832 | 0.097 | 0.194 | 0.499 | 0.624 |
| log(Mass) |  |  | 0.832 | 0.089 | 9.385 | 2.35E-08 |
| log(MeanC) ~ log(SVL) + | 34 | 0.26 | 0.724 | 0.215 | 3.364 | 0.002 |
| Alkaloid presence |  |  | 0.063 | 0.134 | 0.469 | 0.642 |
| log(MeanC) ~ log(cell size) + | 30 | 0.33 | 0.874 | 0.231 | 3.781 | 0.0008 |
| Alkaloid presence |  |  | 0.009 | 0.126 | 0.075 | 0.941 |
| log(MeanC) ~ log(Scope) + | 21 | 0.474 | -0.302 | 0.158 | -1.911 | 0.0729 |
| log(Mass) + |  |  | 0.477 | 0.137 | 3.474 | 0.0029 |
| Alkaloid presence |  |  | -0.015 | 0.129 | -0.118 | 0.907 |
| log(MeanC) ~ log(AMR) + | 21 | 0.491 | -0.375 | 0.181 | -2.073 | 0.053 |
| log(Mass) + |  |  | 0.535 | 0.153 | 3.489 | 0.0028 |
| Alkaloid presence |  |  | -0.007 | 0.127 | -0.06 | 0.954 |
| log(MeanC) ~ log(Number of oocytes) + | 18 | 0.316 | 0.333 | 0.11 | 3.017 | 0.009 |
| Alkaloid presence |  |  | 0.389 | 0.229 | 1.698 | 0.11 |
| log(MeanC) ~ log(Mean_clutch_size) + | 29 | 0.148 | 0.201 | 0.088 | 2.272 | 0.0315 |
| Alkaloid presence |  |  | 0.229 | 0.155 | 1.479 | 0.151 |
